# A brainstem map of orofacial rhythms

**DOI:** 10.1101/2025.01.27.635041

**Authors:** Heet Kaku, Liu D. Liu, Runbo Gao, Steven West, Song-Mao Liao, Arseny Finkelstein, David Kleinfeld, Alyse Thomas, Sri Laasya Tipparaju, Karel Svoboda, Nuo Li

## Abstract

Rhythmic orofacial movements, such as eating, drinking, or vocalization, are controlled by distinct premotor oscillator networks in the brainstem. Orofacial movements must be coordinated with rhythmic breathing to avoid aspiration and because they share muscles. Understanding how brainstem circuits coordinate rhythmic motor programs requires neurophysiological measurements in behaving animals. We used Neuropixels probe recordings to map brainstem neural activity related to breathing, licking, and swallowing in mice drinking water. Breathing and licking rhythms were tightly coordinated and phase-locked, whereas intermittent swallowing paused breathing and licking. Multiple clusters of neurons, each recruited during different orofacial rhythms, delineated a lingual premotor network in the intermediate nucleus of the reticular formation (IRN). Local optogenetic perturbation experiments identified a region in the IRN where constant stimulation can drive sustained rhythmic licking, consistent with a central pattern generator for licking. Stimulation to artificially induce licking showed that coupled brainstem oscillators autonomously coordinated licking and breathing. The brainstem oscillators were further patterned by descending inputs at moments of licking initiation. Our results reveal the logic governing interactions of orofacial rhythms during behavior and outline their neural circuit dynamics, providing a model for dissecting multi-oscillator systems controlling rhythmic motor programs.

## Introduction

Eating and drinking involve temporally patterned orofacial movements, such as the rhythmic tongue protrusions when consuming liquids or the alternating jaw movements of chewing solid food ^1^. These rhythmic orofacial movements are controlled by distinct central pattern generators in the brainstem ^2–9^. Each premotor network oscillates at its own distinct frequency and different rhythms must be coordinated with breathing ^10–12^. Discoordination of orofacial rhythms leads to aspiration pneumonia, a major cause of death among the elderly ^13^ and in patients suffering from neurodegenerative disorders such as Alzheimer’s disease and Parkinson’s disease ^14–16^. It remains poorly understood how brainstem circuits pattern distinct orofacial rhythms relative to breathing and how these dynamics interact with descending control.

Known examples of orofacial coordination include rhythmic whisking and vocalization in rodents ^17–21^, which have been linked to medullary oscillators that interact with the breathing oscillator, the preBötzinger complex ^22,23^. However, the brainstem circuits generating the rhythms underlying eating and drinking have not been comprehensively mapped. Understanding the coordination of these orofacial rhythms requires identification of brainstem premotor networks and measuring their dynamics during behavior. Yet, neurophysiological recording in the brainstem of awake behaving animal models remains technically challenging.

Here we examine the coordination of breathing, licking, and swallowing in mice drinking water. Mice gather water by stereotyped rhythmic licking at 7 Hz ^5^ while breathing occurs across a range of frequencies between 1 and 10 Hz ^18^. Measurements in rats show that licking is time-locked to inspiration ^24^ but the directionality of licking-breathing coupling is obscure. Bouts of licking accumulate liquid in the oral cavity to trigger swallowing, which in turn inhibits beathing ^11,25,26^, similar to drinking in humans ^27^. The interplay between these rhythms enables investigation of how different orofacial rhythms are temporally organized into coherent ingestive behavior.

Movement of the jaw and tongue muscles for licking is controlled by motor neurons in the trigeminal nucleus (5N) and hypoglossal nucleus (12N), respectively ^5^. Swallowing is controlled by the nucleus ambiguus (10N) that innervates the muscles of the pharynx, larynx, and esophagus ^6^. Upstream of these primary motor nuclei, the location of brainstem premotor networks generating and coordinating orofacial rhythms are not well understood. Rhythmic licking is thought to be controlled by a neural oscillator in the intermediate reticular nucleus (IRN) of the medulla ^5,9,28^, and a group of *Phox2b^+^* neurons in the IRN has been reported to elicit licking ^9^. But IRN is a large structure with diverse functions ^18,19,21,26,29–31^ and functional nuclei within IRN have not been parsed.

We used extracellular recordings with Neuropixels probes to create a brainstem-wide activity map of behaving mice. Breathing, licking, and swallowing were tightly coordinated: breath and lick timing reciprocally influenced each other, while intermittent swallowing paused both breathing and licking rhythms. Distinct clusters of neurons exhibited activity time-locked to breathing, licking, and swallowing. Licking-related activity delineated a lingual premotor network in the IRN. Optogenetic stimulation mapping further identified a distinct region with rhythm-generation properties capable of driving rhythmic licking. By artificially activating the licking circuits to induce licking, we found that brainstem oscillators autonomously coordinated structured breathing and licking, with simultaneous recordings revealing functional coupling between the two oscillators. We also uncovered coordination not accounted for by the brainstem circuits: when mice initiated licking, ongoing breathing was pre-emptively adjusted to accommodate coupled licking-breathing oscillations and allow rapid movement initiation, revealing descending control of orofacial movements. These results reveal the logic governing orofacial rhythms of ingestive behavior and outline their brainstem circuit dynamics, providing a foundation for understanding multi-oscillator systems controlling rhythmic behavior.

## Results

### Coordination of breathing, licking, and swallowing

We measured breathing, licking, and swallowing in mice drinking water from a spout to examine how the three orofacial rhythms are temporally organized. Mice performed a directional licking task in which they licked to one of nine targets for a water reward presented on a lickspout ^32^ (**Figure 1a**). Orofacial movements were recorded with high-speed videography while breathing was recorded with an airflow meter (**Figure 1b**) (Methods). We measured licking by tracking the movement of jaw and tongue using DeepLabCut ^33^ (Methods; **Supplemental Movie 1**; tongue protrusions were defined as the moments when the tongue appeared in the video, and individual licks were defined as the onset of each tongue protrusion). Swallowing was accompanied by a pause in breathing ^24,25^ (**Figure 1c**). We used a heuristic algorithm to identify swallowing events based on the beathing pattern (Methods; **Extended Data Figure 1a**). Although we used breathing traces to classify swallowing, licking was also paused during the swallowing events, suggesting that the classified events were genuine swallows (**Extended Data Figure 1b-c**; also see single-unit activity below).

**Figure 1.**
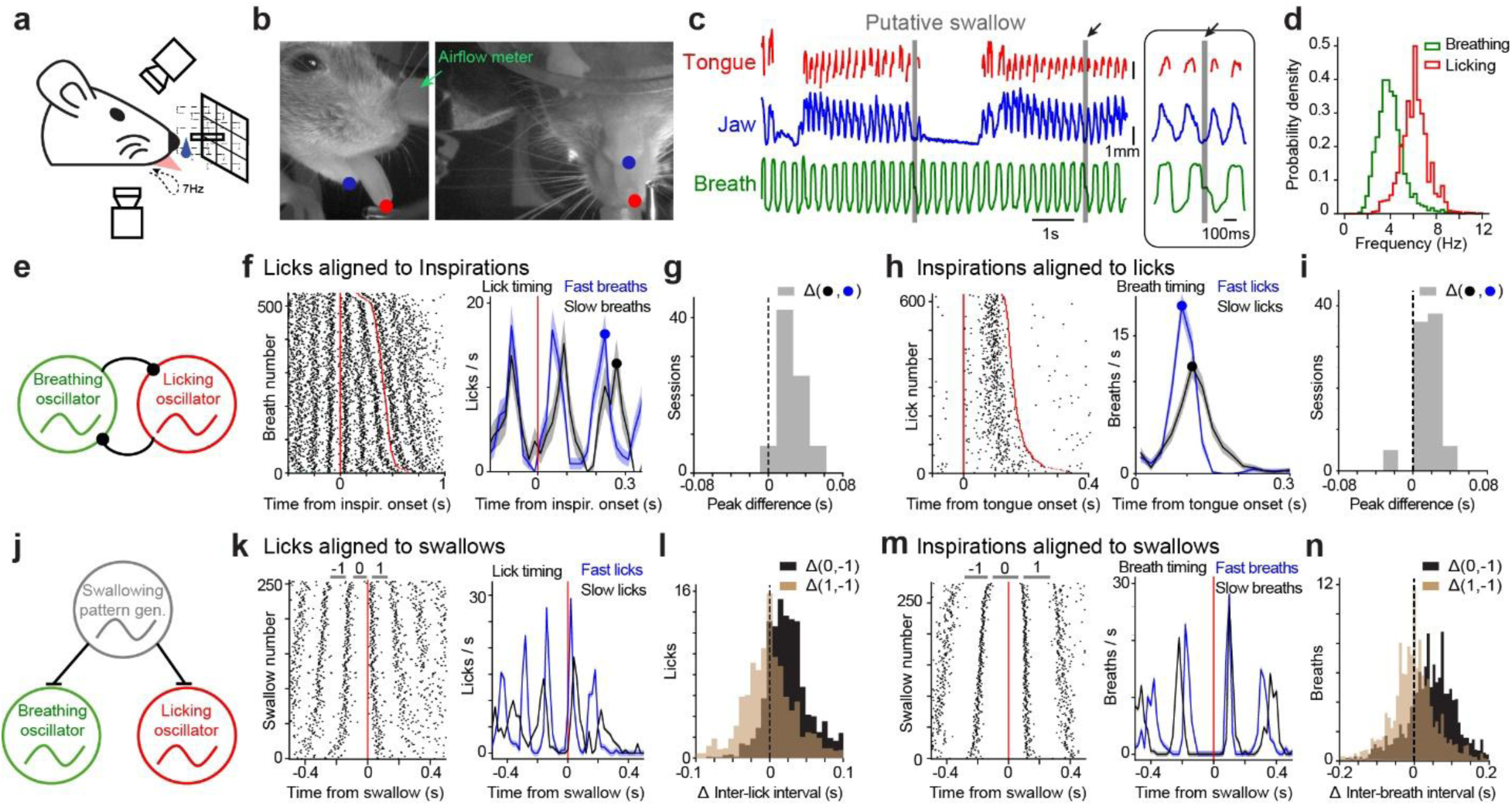
Coordination of orofacial rhythms in drinking mice. **a**) Multi-directional licking task. On a given trial, the lick spout moved to one of nine possible positions and the animals licked the spout to retrieve a water reward. High-speed videography was made from the bottom view and side view cameras. **b**) Example bottom view and side view video frames with the tip of the tongue and the jaw labelled. **c**) Example traces of tongue protrusions (red), jaw movement (blue), breathing (green), and swallowing (gray). Inset, zoomed in view of the example swallow indicated by the arrow. **d**) Histograms of licking and breathing frequencies. **e**) Schematic illustrating bidirectional coupling of the breathing and licking oscillators. **f**) *Left*, licks aligned to breaths in an example session. The timing of each lick is represented by a dot plotted against inspiration onset, red lines. The inspirations are ranked by the inter-breath interval where the shortest breath is at the top. *Right*, lick times for trials with two licks within a breathing cycle, sorted by breath frequency. Blue, the top half of the trials; black, the bottom half of the trials. The licks precede inspirations with a fixed latency, and the peak time for the second lick is delayed for slower breath frequency (blue vs. black dot). **g**) The distribution of lick time difference between top and bottom half of trials (blue vs. black dot in **f**) for all sessions. The distribution is shifted to positive. **h-i**) Breaths aligned to licking; same as panels **f-g**. **j**) Schematic of the swallowing pattern generator inhibiting the breathing and licking oscillators. **k**) *Left*, licks aligned to swallows in an example mouse. The timing of each lick is represented by a dot plotted against swallow onset, red lines. The licks are ranked by the inter-lick interval where the faster licking frequency is at the top. *Right*, lick times of the top 1/3 of the trials (blue) and the bottom 1/3 of the trials (black). **l**) The change in inter-lick intervals for bottom 1/3 of the trials. Lick cycles ‘-1’, ‘0’, and ‘1’ are labeled in **k**. Δ(1, -1), difference between the inter-lick intervals before and after swallowing (cycle ‘1’ minus ‘-1’). Δ(0, -1), difference between the inter-lick intervals before and during swallowing (cycle ‘0’ minus ‘-1’). Inter-lick interval is prolonged during swallowing but recovers after swallowing. **m-n**) Inspirations aligned to swallowing; same panels as **k-l**.

Licking occurred at 6.23 ± 2 Hz (mean ± std, 95% range: 3.59 - 8.91 Hz) while breathing occurred at lower frequencies (4.02 ± 1.24 Hz, 95% range: 2.12 - 7.17 Hz; **Figure 1d**). Despite their different frequencies, the two rhythms were tightly coordinated. To visualize this coordination, we plotted the timing of licks relative to inspiration sorted by breathing frequency (**Figure 1f**). Inspiration always occurred between licks and thus never coincided with licking. Licking preceded inspiration at a fixed latency (**Figure 1f**, the 1st red line) and the same lick-breath temporal relationship followed in the next inspiration across the entire range of breathing frequencies (**Figure 1f**, the 2^nd^ red line). For slower breathing frequencies, the licking frequency between the two inspirations slowed down in anticipation of the next breath such that the next lick always preceded the next inspiration at a fixed latency. To quantify this anticipatory adjustment, we calculated the lick timing preceding the next inspiration (**Figure 1f** right panel, blue vs. black dot, comparing top and bottom trials for trials with two licks between breath). In all mice, the lick timing was significantly delayed preceding a slower breath compared to a fast breath (**Figure 1g**, p<0.001, one sample t-test, two-tailed). These results indicate that breathing influences the licking rhythm.

The range of licking frequencies was faster than breathing frequencies (**Figure 1d**). During slow breathing, when the next lick could not be delayed further to accommodate the next breath, licking frequency abruptly increased to add additional licks within the breathing cycle to maintain the fixed lick latency preceding the next inspiration (**Figure 1f**, compare trials with 1 vs. 2 vs. 3 vs. 4 licks between breaths). Thus, licking rhythm shifted between different modes of oscillation depending on the breathing frequency.

We next examined whether the breathing rhythm was also influenced by licking. If the breathing oscillator is free-running and unidirectionally adjusts the timing of licking rhythm, inspiration timing should be insensitive to future lick timing (**Extended Data Figure 1d-f**). We visualized the timing of inspiration relative to licking in a frequency-ordered plot sorted by licking frequency (**Figure 1h**). Although licking spanned a limited range of frequencies, inspirations following licking were significantly delayed for slower licking frequencies to anticipate the next lick (**Figure 1i**, p<0.001, one sample t-test, two-tailed). This pattern of lick-breath temporal relationship also cannot be produced by a free-running licking oscillator unidirectionally setting the timing of breathing rhythm (**Extended Data Figure 1g-i**). Independent analysis of breathing and licking in freely moving rats yielded similar temporal relationship (**Extended Data Figure 2**). Together, the interdependence of licking and breathing rhythms suggest that their underlying neural oscillators are bidirectionally coupled.

Swallowing inhibited both breathing and licking. We examined its influence on licking and breathing rhythms by aligning the timing of inspiration and licking to swallowing, ordered by their frequencies before the swallow (**Figure 1k-n**). Inspirations and licks followed swallowing at relatively fixed latencies despite different breath and lick timing before the swallow (**Figure 1k and m**), suggesting that ongoing rhythms were interrupted by swallowing. We tested whether swallowing altered the frequency of licking by comparing the inter-lick-intervals around the swallowing event. The inter-lick-interval before and after swallowing was preserved (**Figure 1l**, P=0.649, one-sampled t-test, two-tailed), whereas the lick cycle in which swallowing occurred was significantly prolonged by 33.3 ms (**Figure 1l**, P<0.001, one-sampled t-test, two-tailed). The same pattern was also observed for breathing rhythm (**Figure 1n**; P=0.225 for inter-breath-interval before vs. after swallowing; P<0.001 for inter-breath-interval in which swallowing occurred, one-sample t-test, two-tailed, prolonged by 48.0 ms). These results indicate that swallowing paused the ongoing licking and breathing oscillators without changing their frequencies.

Together, these results identify the logic governing the coordination of breathing, licking, and swallowing over rapid timescales (<100 ms) during drinking behavior.

### Brainstem activity maps of orofacial rhythms

Breathing is controlled by a central pattern generator in the preBötzinger complex ^22^. The relevant effector motor neurons for licking and swallowing are known: tongue protrusion and jaw movements are controlled by the hypoglossal nucleus (12N) and trigeminal nucleus (5N) respectively ^5^, whereas swallowing is controlled by the nucleus ambiguus (10N) ^6^. But the locations of brainstem premotor networks generating and coordinating licking and swallowing have not been mapped.

We used Neuropixels probes to record brainstem activity related to breathing, licking, and swallowing in behaving mice. We inserted 1-2 Neuropixels 1.0 probes at a time to target the brainstem (**Figure 2a-c**). Across 339 insertions from 31 animals, we sampled activity bilaterally across the hindbrain, covering the entire medulla and a large portion of the pons (**Figure 2d**). After spike sorting with Kilosort 2.5 ^34^ and stringent quality control metrics ^35^ (Methods), we obtained 20,300 single units (60 ± 22 units per probe, mean ± std across recordings). The unit locations were reconstructed within the Allen Mouse Common Coordinate Framework (CCFv3) using a combination of histological information and electrophysiological landmarks (**Extended Data Figure 3**; Methods) ^36^. This allowed us to map activity patterns within the brainstem and its anatomical compartments (**Figure 2e**) ^37^. We confirmed that major results were insensitive to the quality of spike sorting, including activity modulation by licking (**Extended Data Figure 4**).

**Figure 2.**
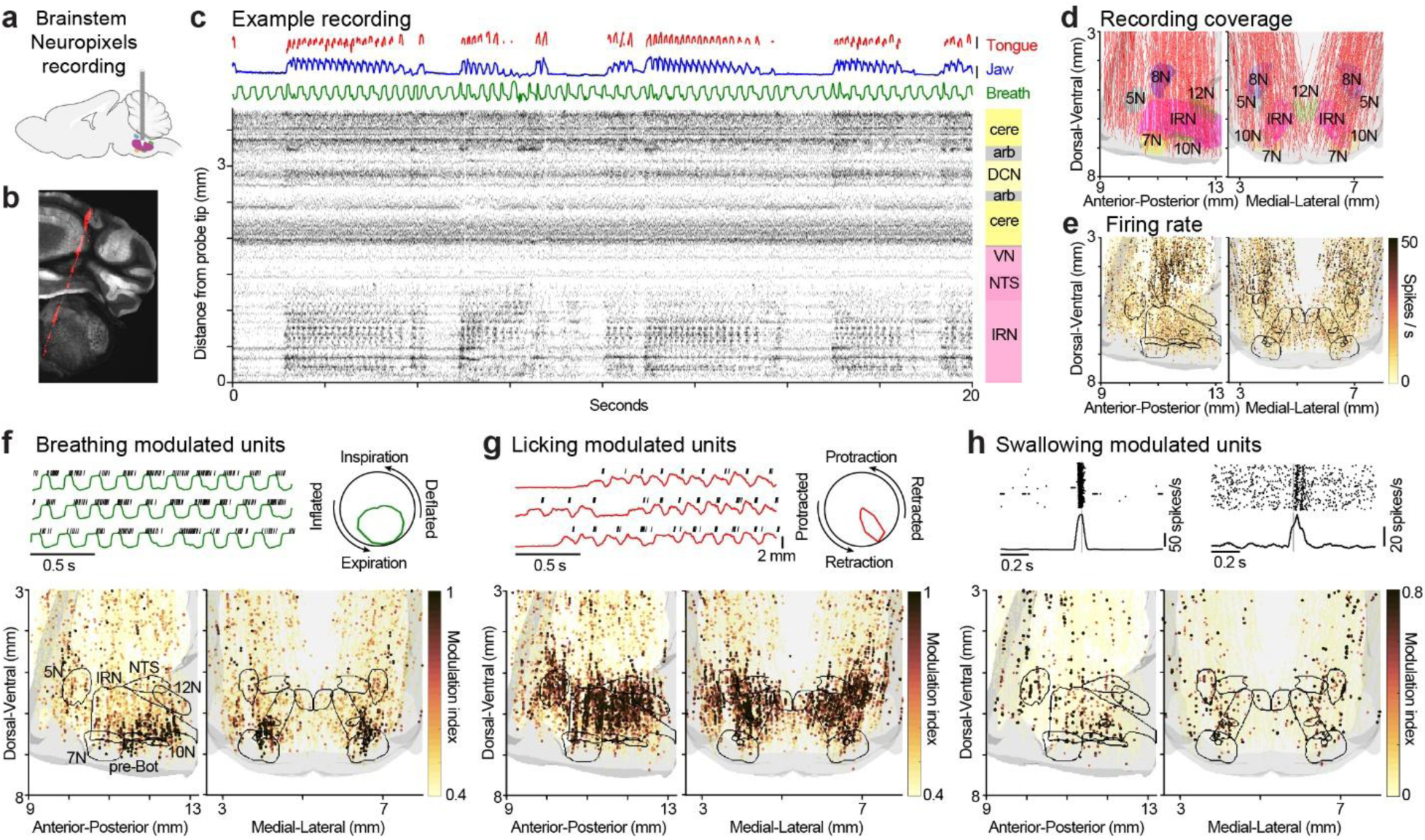
A brainstem activity map during orofacial behaviors. **a**) Neuropixels recordings in the brainstem. **b**) Example warped image to the Allen anatomical template brain and example probe insertion. The gray is autofluorescence and red is DiI fluorescence from the probe track. **c**) Example tongue, jaw, and airflow tracking with spikes along a probe insertion. Scale bars, 2 mm. Spikes along the probe are averaged in time bins of 2.5 ms and depth of 20 µm. The example probe insertion shows licking-related activity in the IRN. The ARA annotation of the localized electrode sites is shown on the right. **d**) Coverage of the recordings in CCF with probe tracks labelled in red and brainstem nuclei. 5N: Trigeminal Nucleus. 7N: Facial Nucleus. 8N: Vestibular Nucleus. 10N: Nucleus Ambiguus. 12N: Hypoglossal Nucleus. IRN: Intermediate Reticular Nucleus. **e**) Localized units as dots and spike rate indicated by the color. The outlines denote the motor nuclei in **d**. **f**) *Top*, a breathing-related unit with three trials of airflow traces in green and spike times indicated by ticks on top. The tuning curve of the unit is shown on the right. *Bottom*, activity map of breathing. The modulation index for breathing of every unit is indicated by the darkness of the color. Regions with high density of black dots indicate highly tuned units to breathing. pre-Bot, the preBötzinger complex. **g**) *Top*, a licking-related unit with three trials of jaw movement traces in red and spike times indicated by ticks on top. The tuning curve for the unit is shown on the right. *Bottom*, activity map of licking. Same as **f**. **h**) *Top*, rasters and PSTHs of two swallowing-related units with spike times aligned to swallowing onset (gray line). *Bottom*, activity map of swallowing. Same as **f**.

We correlated the firing of individual neurons to orofacial rhythms. A subset of units exhibited rhythmic activity phase-locked to breathing (**Figure 2f**, top). A different subset of the units were active during licking and they fired at specific phases of rhythmic licking (**Figure 2g**, top). We quantified breathing- and licking-related activity by calculating firing rate modulation across breathing or licking phase (**Extended Data Figure 5a** and **c**; permutation test, P < 0.001, and modulation index>0.6; Methods). Individual units exhibited significant phase tuning to either licking (3551/20300), or breathing (1015/20300), but rarely to both (182/20300). Additional brainstem units showed transient activity during swallowing (**Figure 2h**, top). We quantified swallowing-related activity by significant firing rate modulation during swallowing events (100 ms) compared to 200 ms time windows prior and after (P<0.01, bootstrap, and modulation index>0.3; **Extended Data Figure 5e**). Most units were modulated by swallowing only (427/18485), with a smaller group modulated by both licking and swallowing (175/18485) (**Extended Data Figure 5f**). Another small subset of units with swallowing-related activity were also modulated by breathing (55/18485). The largely non-overlapping tunings suggest distinct brainstem networks for specific behaviors.

Neurons with activity related to breathing, licking, and swallowing were clustered in distinct locations (**Figure 2f-h**, bottom). Units encoding breathing were near the preBötzinger complex ^22^, as well as posterior ventral IRN that receives inputs from the preBötzinger complex to coordinate whisking and breathing ^18^ (**Figure 2f**). The regions containing licking-related activity were remarkably broad, including the tongue- and jaw-related motor nuclei (hypoglossal nucleus, 12N; trigeminal nucleus, 5N) as well as extensive portions of the IRN and parts of pons (**Figure 2g**). Units with swallowing-related activity were clustered in the trigeminal nucleus (5N), lateral to the nucleus of the solitary tract (NTS), and above the nucleus ambiguus (10N) (**Figure 2h and Extended Data Figure 6a-d**). This swallowing activity pattern aligns with previous findings, where a dorsal premotor network near the NTS triggers swallowing, and a ventral premotor network above the 10N distributes swallowing signals to motoneurons ^6^. Unexpectedly, additional clusters of swallowing-related neurons were found in the inferior colliculus (IC) and nucleus of the lateral lemniscus (NLL) (**Extended Data Figure 6a**), possibly elicited by the sound of swallowing or an efference copy signal ^38^ as suggested by their response latencies (**Extended Data Figure 6e**).

We next analyzed the neural coding of the brainstem units, focusing on encoding of licking and breathing. Tracking of tongue and jaw movements together with breathing airflow provided reduced measures of these orofacial rhythms. We used a generalized linear model (GLM) to predict the spike rates of individual units using combinations of these tracked features (**Extended Data Figure 7a-b**). The GLM found separate clusters of units predicted by either licking or breathing movement (**Extended Data Figure 7c-d**), consistent with activity modulations by licking and breathing (**Figure 2f-g**). For units with breathing-related activity, units in the preBötzinger complex preferentially fired at inspiration phase of breathing (**Extended Data Figure 8a-c**), consistent with their role in driving inspiration ^22,23^. Units with licking-related activity were mostly active when the tongue was in the fully protruded state (**Extended Data Figure 8d-f**)^39^, presumably to support the fully protracted state of the genioglossus muscle.

We tested whether the licking- and breathing-related neurons encode movement kinematics beyond rhythmicity. To capture movement kinematics in the videos beyond the tracked features, we trained a deep convolutional autoencoder (CAE) to compress the high dimensional videos to a 32-dimensional embedding ^40^ (**Extended Data Figure 7e-f**). We then used a GLM to predict the spike rates of individual units from the 32-dimensional embedding features. The CAE predicted a larger fraction of activity variance than the tracked features for most units across the brainstem (**Extended Data Figure 7g;** CAE vs. features, P<0.001, Wilcoxon signed rank test). Interestingly, the only exceptions were units in the preBötzinger complex where activity prediction using CAE led to worse predictions (CAE vs. features, P<0.001, Wilcoxon signed rank test; **Extended Data Figure 7g-h**). This suggests that a 1-dimensional signal primarily capturing rhythmic movement was sufficient to explain the activity of the breathing oscillator while majority of brainstem regions exhibited rich encoding of movement parameters.

Finally, we tested whether the licking-related units were sensitive to tongue kinematics. Mice directed their tongue to multiple directions in the directional licking task. We grouped individual tongue protractions by direction and analyzed individual neuron firing rates as a function lick direction (**Extended Data Figure 8g-h**). Overall, licking-related neurons showed selectivity for ipsilateral licking (**Extended Data Figure 8i**), consistent with drive to contract intrinsic tongue muscles to perturb the tongue into ipsilateral space ^41,42^. However, individual units within each hemisphere exhibited diverse direction selectivity, including units selective for the center lick direction (**Extended Data Figure 8h**).

Together, these brainstem activity maps reveal distinct clusters of neurons for breathing, licking, and swallowing with rich encoding of movement parameters.

### A licking premotor network with autonomous rhythm-generation properties

The spatial distribution of licking-related units in the brainstem was unexpectedly broad (**Figure 2g**), encompassing large territories of medulla and parts of pons beyond the tongue-jaw motor nuclei (hypoglossal nucleus, 12N, and trigeminal nucleus, 5N). This distributed activity may reflect concerted recruitment of tongue, jaw, and facial muscles involved during licking. We next sought to delineate the licking premotor network.

Rhythmic licking is thought to be driven by a central pattern generator in the brainstem ^5,9,28^. Thus, the premotor network should innervate the relevant motor nuclei involved in licking, be innervated by descending inputs from the higher-order licking motor centers, and have rhythm-generation properties.

First, we analyzed our electrophysiology recordings in relation to brainstem premotor neurons innervating the licking muscles. Licking involves movement of the jaw by the masseter muscle and protraction of the tongue by the genioglossus muscle. We mapped premotor neurons of 12N and 5N labeled using retrograde rabies virus transsynaptic tracing from the masseter and genioglossus muscles to the CCF (**Figure 3a** and **Extended Data Figure 9a-b**; data from ^43^; AAV2-retro-Cre in the muscles, Cre-dependent G and TVA along with RV-ΔG in the motor nucleus). Premotor neurons for the masseter and genioglossus muscles occupied largely distinct regions of the pons and medulla, with some overlap in the dorsal IRN (**Figure 3a**). Units synchronized to licking coincided with the tongue-jaw premotor regions (**Figure 3b**). Units inside these regions showed significantly higher licking modulation compared to position-shuffled populations (P < 0.001 Wilcoxon rank sum test). The dynamics within these premotor regions may organize the temporal structures of licking and breathing while activity in the motor nuclei 5N and 12N may reflect motor drive to tongue-jaw muscles.

**Figure 3.**
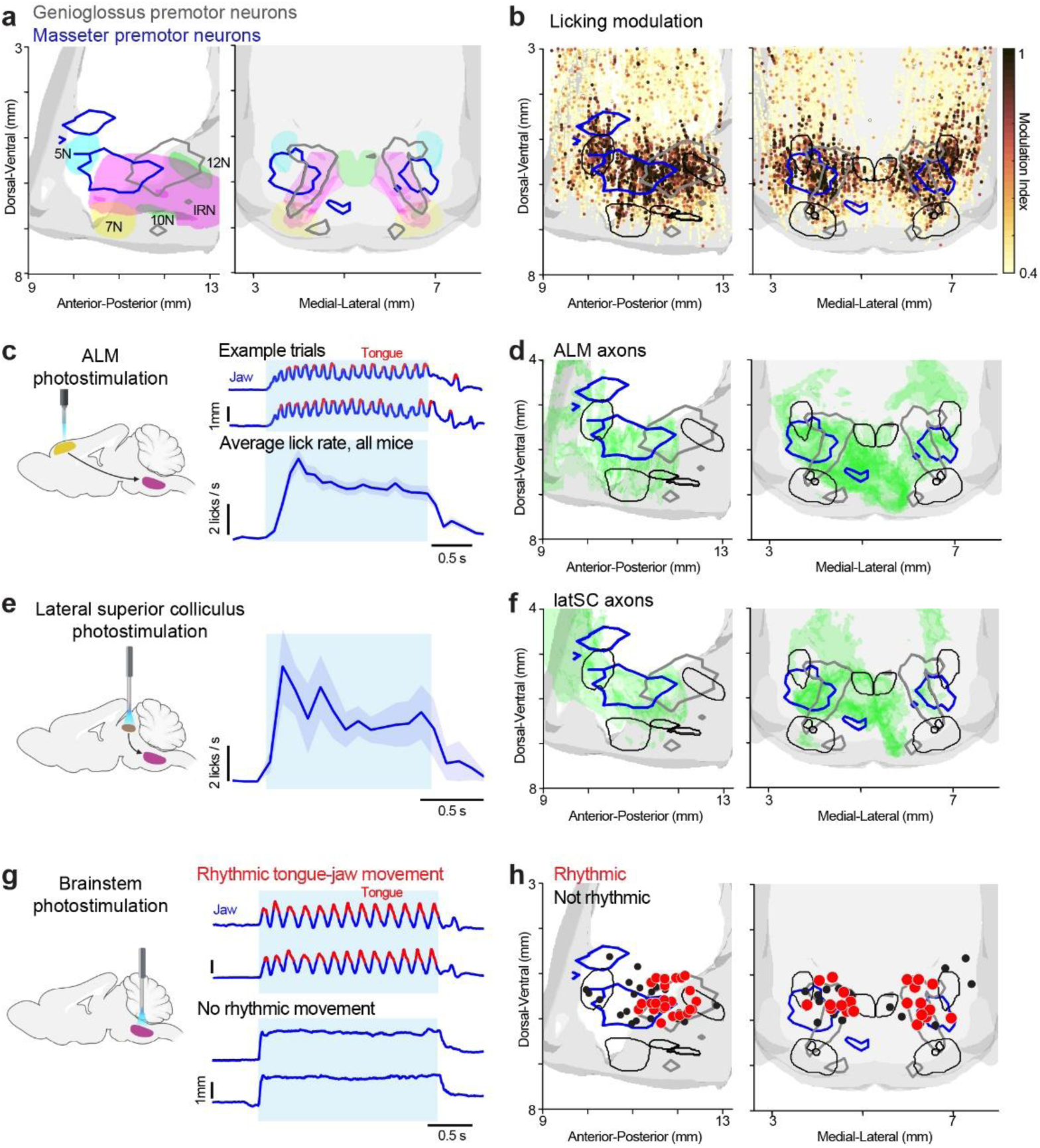
Brainstem premotor network for licking. **a**) Boundaries of the premotor neurons for the masseter (blue) and genioglossus (grey) muscles. Colored regions indicate the motor nuclei. 5N: Trigeminal Nucleus. 7N: Facial Nucleus. 10N: Nucleus Ambiguus. 12N: Hypoglossal Nucleus. IRN: Intermediate Reticular Nucleus. **b**) Activity map of licking. Same as Fig 2g, with the boundaries of tongue-jaw premotor regions and motor nuclei overlaid. **c**) Photostimulation of ALM evokes sustained rhythmic licking. Example jaw movement (blue) and tongue protrusions (red segments) for example photostimulation trials and average lick rate during photostimulation (cyan). Mean ± SEM across mice. n=11 mice. **d**) Fluorescence from ALM axon projections (green), with the boundaries of tongue-jaw premotor regions and motor nuclei overlaid. **e**) Photostimulation of the lateral SC (latSC) evokes sustained rhythmic licking. Same as **c**. n=3 mice. **f**) Fluorescence from latSC axon projections (green), with the boundaries of tongue-jaw premotor regions and motor nuclei overlaid. **g**) Photostimulation of the brainstem. Example jaw movement (blue) and tongue protrusions (red segments) for a photostimulation site where sustained rhythmic licking was evoked and a photostimulation site where rhythmic licking could not be evoked. Cyan, photostimulation epoch. **h**) Spatial map of photostimulation sites. Each dot shows one optical fiber implant, n=46 photostimulation sites, 27 mice. Red dots, photostimulation sites where rhythmic licking was elicited. Black dots, photostimulation sites where rhythmic licking was not elicited.

We next analyzed the innervation patterns of higher-order licking centers in the brainstem in relation to the licking activity map. The anterior lateral motor cortex (ALM) ^42,44–47^ and the lateral superior colliculus (latSC) ^48–51^ are known to elicit licking. To confirm that these descending inputs can elicit rhythmic licking, we expressed ChR2 in ALM layer 5 pyramidal neurons (Sim1_KJ18-cre x Ai32 or Tlx_PL56-Cre x Ai32 mice) or in the latSC (ChR2 virus) and photostimulated these regions (Methods; unilateral photostimulation). Photostimulation of ALM or latSC (20 or 40 Hz pulse trains for 2 or 1.3 s) elicited sustained rhythmic licking at the naturalistic 7 Hz frequency (**Figure 3c and e**), consistent with these descending inputs activating a neural oscillator in the brainstem to drive licking. Next, we measured mesoscale projections using anterograde tracers injected in ALM or latSC (**Extended Data Figure 9c-d**). ALM and latSC axons targeted the contralateral brainstem and showed similar innervation patterns, which engulfed the tongue-jaw premotor regions enriched with licking activity (compare **Figure 3b, d, and f**). Thus, the tongue-jaw premotor regions with licking activity outline the premotor network for licking.

Finally, we tested if the outlined licking premotor network can generate rhythms autonomously. A group of *Phox2b^+^* neurons in the IRN has been reported to elicit licking ^9^, but *Phox2b* is a neurodevelopmental transcription factor expressed throughout the brainstem ^52,53^ and the site capable of eliciting rhythmic licking has not been mapped. We therefore expressed ChR2 or ChRmine in *Phox2b^+^* neurons and photostimulated different locations across the identified licking premotor network (**Figure 3g**; Methods; *Phox2b-cre x Ai32* mice or cre-dependent ChR2 or ChRmine viruses in *Phox2b-cre* mice; **Extended Data Figure 9e-f**). We used constant light stimulus to identify regions with rhythm-generation properties. Photostimulation sites in the anterior regions of the licking premotor network elicited jaw movements. But these movements were not rhythmic: the jaw was locked to a fixed position during photostimulation and there was little or no tongue protrusion (**Figure 3g-h and Extended Data Figure 10**; see **Supplemental movie 2**). A few photostimulation sites near 5N elicited a transient tongue protrusion. But unlike rhythmic licking, the tongue stopped after one or two protrusions and the jaw was locked (**Extended Data Figure 10**; see **Supplemental movie 3**). In contrast, photostimulating a posterior-dorsal region of IRN and the adjacent region in PARN (anterior-lateral to 12N) elicited rhythmic jaw movement and tongue protrusions (**Figure 3g-h** and **Extended Data Figure 10**; here on referred to ‘posterior-dorsal IRN/PARN’, or IRt/PCRt by Paxino convention ^54^). These rhythmic movements were sustained for the entire photostimulation period and ceased upon termination of the light (see **Supplemental movie 4**). This region is thus capable of generating licking rhythms under constant input.

These results outline a brainstem premotor network for licking and a distinct region within capable of autonomously generating rhythms to drive licking. The posterior-dorsal IRN/PARN is likely a core part of the licking oscillator. Notably, the licking premotor network identified with electrophysiology is larger than the posterior-dorsal IRN/PARN region that triggers rhythmic licking (**Figure 3b vs. h**). The effect of photostimulation could spread beyond the targeted neurons.

### Coupled brainstem oscillators autonomously coordinate licking and breathing

We next sought to understand how licking and breathing rhythms are organized by their premotor circuits given their tight cycle-by-cycle coordination during drinking behavior (**Figure 1e-i**). Breathing and licking could be patterned autonomously by coupled brainstem oscillators. Alternatively, conflicts between different oscillations could be resolved downstream of rhythmogenesis through gating of motor nuclei. Finally, it is also possible that different rhythms are coordinated by descending control from higher-order motor centers.

We first tested if licking and breathing could be autonomously coordinated under conditions where the licking oscillator is driven artificially. We used ALM photostimulation because it robustly activates the licking oscillator (**Figure 3c**) and ALM projections avoid the breathing oscillator (**Figure 3d and Extended Data Figure 9c**). We photostimulated ChR2 in ALM layer 5 neurons (**Figure 4a**; Methods; Sim1_KJ18-Cre x Ai32 or Tlx_PL56-Cre x Ai32 mice, 20 Hz pulse trains for 2 s). We tested mice in a cued licking task, with the goal to compare artificially induced licking to voluntary licking. Mice licked for water from a fixed lickspout after a randomly timed Go cue (Methods). Licking and breathing maintained a faithful temporal relationship during voluntary licking (**Figure 4b-c**). On a random subset of trials, the Go cue was substituted with ALM photostimulation and water reward was withheld on the photostimulation trials to avoid water-induced licking. Photostimulation of ALM induced sustained rhythmic licking (**Extended Data Figure 11a**)^42^. In a subset of mice, ALM photostimulation also induced an increase in breathing rate (**Extended Data Figure 11b**). Notably, the same lick-breath temporal relationship was recapitulated during artificially induced licking. Lick timing anticipated the next breath and proceeded inspiration at a fixed latency across different breathing frequencies (**Figure 4b**). Conversely, inspiration timing was influenced by future lick timing (**Figure 4c**). The 20 Hz stimulation disrupted natural patterns of activity in ALM, likely hindering its ability to coordinate licking with breathing on a cycle-by-cycle basis. This suggests that the brainstem circuits may autonomously coordinate licking and breathing.

**Figure 4.**
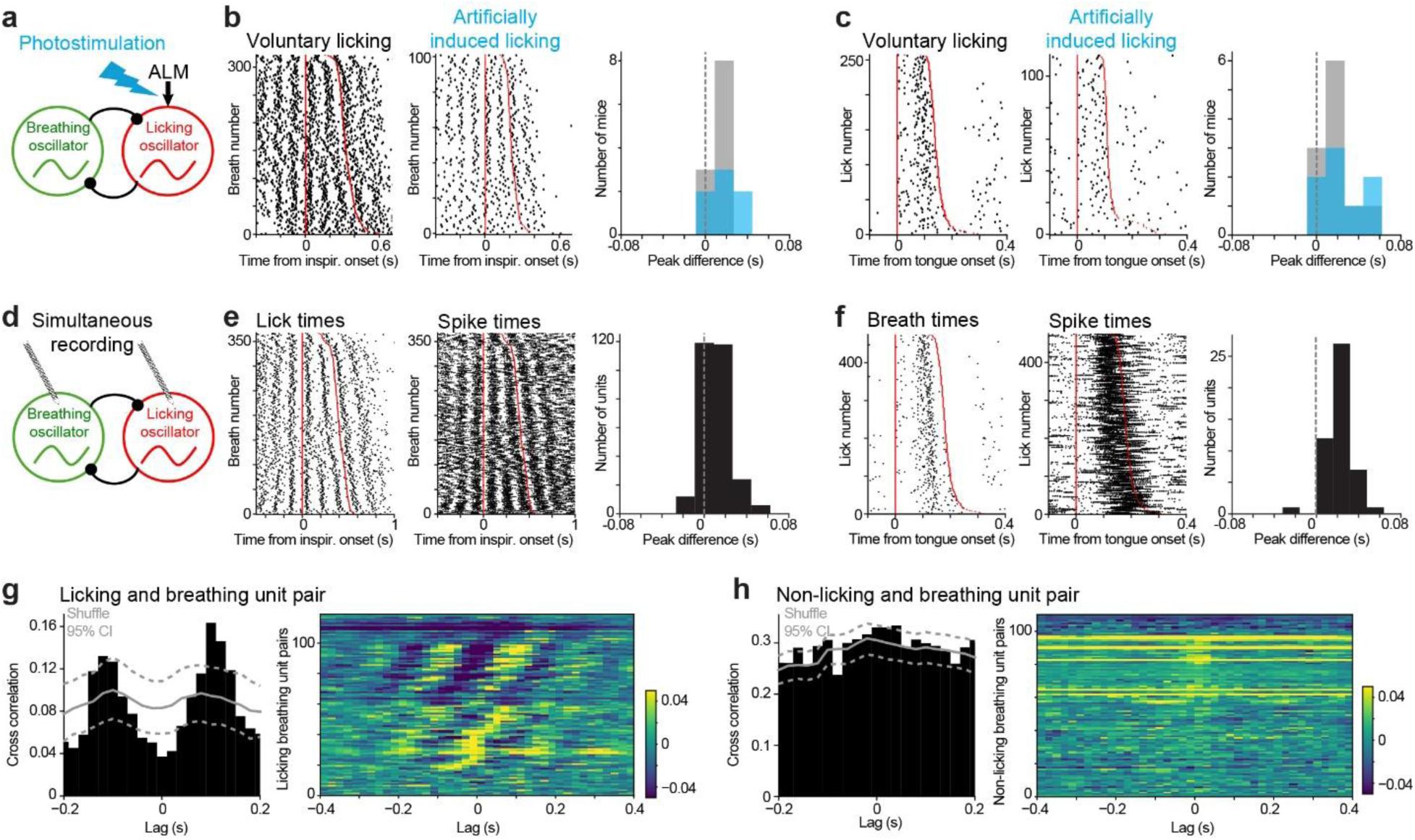
Coupled brainstem oscillators coordinate licking and breathing. **a**) ChR2 stimulation of ALM to activate the licking oscillator and elicit licking. **b**) *Left*, lick-breath temporal relationship during voluntary licking and artificially induced licking. Data from an example session. *Right*, the distribution of lick time difference for slow vs. fast breath across all mice. Same as Fig 1f-g. The same lick-breath temporal relationship is preserved during artificially induced licking. **c**) Breaths aligned to lick; same as Fig 1h-i. **d**) Simultaneous recording from the licking and breathing oscillators. **e**) An example unit from the licking oscillator. *Left*, lick timing and spike timing aligned to breaths. *Right*, the distribution of spike time difference for slow vs. fast breath across all sessions. Same as **b**. **f**) An example unit from the breathing oscillator with activity following breathing. Same as **e**. **g**) *Left*, cross-correlogram of an example pair of breathing unit and licking unit plotted in black. The trial-shuffled mean cross-correlogram is plotted as the gray line, 95% confidence interval is indicated by the dashed lines. *Right*, the difference between the measured cross-correlogram and the shuffle for all unit pairs. Unit pairs without significant cross correlation are shown on the bottom; units with positive cross correlations are shown in the middle; units with anticorrelated activity are shown on the top. **h**) *Left*, cross-correlogram of an example pair of breathing unit and non-licking unit. *Right*, population summary for all pairs as in **g**.

To determine if licking and breathing rhythms are directly patterned at the level of rhythmogenesis, we used 2 Neuropixels probes to simultaneously record from the licking and breathing oscillators to examine their dynamics (**Figure 4d**, posterior-dorsal IRN/PARN and preBötzinger complex; 43 recordings in 21 mice; 1-28 licking-related units and 1-12 breathing-related units per recording). Activities in the two regions reflected the coordinated lick-breath temporal structure. Preceding a slower breath, the activity of licking neurons was slowed in anticipation of the next inspiration (**Figure 4e**). Preceding a slower lick, the activity of breathing neurons was slowed in anticipation of the upcoming lick (**Figure 4f**). This indicates that lick and breath timing are already coordinated at the level of rhythmogenesis.

The dependency of licking and breathing oscillator activities on the other rhythm suggests that the two oscillators might be coupled. To look for coupling, we analyzed cross-correlation between simultaneously recorded breathing and licking unit pairs (119 pairs in total, **Extended Data Figure 12a-b**). Activity was significantly correlated for most pairs compared to trial-shuffled controls (**Figure 4g**; 103/119 pairs with significant cross-correlation relative to shuffled control, 43 positively correlated, 60 anticorrelated; Methods). The cross-correlation was temporally symmetric: breathing units lagged licking units for 57 pairs, while other licking units lagged breathing units for 34 pairs, (the remaining 12 pairs had cross-correlation peak at 0). Notably, significant cross-correlation between the unit pairs was observed even when mice were not licking, i.e. when the licking oscillator was inactive (**Extended Data Figure 12c**). This correlated activity suggests reciprocal functional coupling between the licking and breathing oscillators. In contrast, units outside of the licking oscillator exhibited less cross correlation with the breathing units (**Figure 4h**; 54/110 pairs with significant cross correlation; a significantly lower proportion compared to licking-breathing oscillator unit pairs, p<0.0001, Chi-square test).

These data show that licking and breathing rhythms are coordinated at the level of brainstem rhythmogenesis and suggest that coupled licking and breathing oscillators organize the two rhythms over rapid timescales.

### Descending control patterns oscillators at movement initiation

Given the coupled licking-breathing oscillations coordinated by brainstem circuits, does the breathing rhythm dictate mice’s ability to initiate licking? Breathing rhythm spans a slower range of frequencies compared to licking (**Figure 1d**), but the licking-breathing coordination constrains licking to a fixed latency preceding inspiration (**Figure 1f-i**). Are mice obligated to initiate licking only after waiting for a cycle of breath? This is not the case: the latency for cued-evoked licking is much faster than a cycle of breath (250 ms, given an average breathing frequency of 4.0 Hz, **Figure 1d**), and can be faster than 100 ms in some conditions ^55^.

To understand how mice were able to rapidly initiate licking while maintaining the lick-breath temporal relationship, we analyzed movement initiation in the cued licking task. Mice initiated licking 127.6 ± 29.6 ms after the Go cue onset (defined as the onset of tongue appearance in the video, mean ± std across mice, n=11 mice; **Figure 5a-b**). We aligned the phase of breathing to licking onset (**Figure 5c**). Breathing phase was initially random with respect to lick initiation. 115 ± 19 ms before licking onset (mean ± s.e.m., bootstrap), breathing phase began to deviate significantly from random distribution and became entrained to a specific phase (**Figure 5c**; p<0.0001, two sample Kuiper test with Bonferroni correction). This pre-emptive entrainment set the breathing cycle to the expiration state at the time of the first lick. Thus, mice pre-emptively adjusted their breathing rhythm prior to licking initiation.

**Figure 5.**
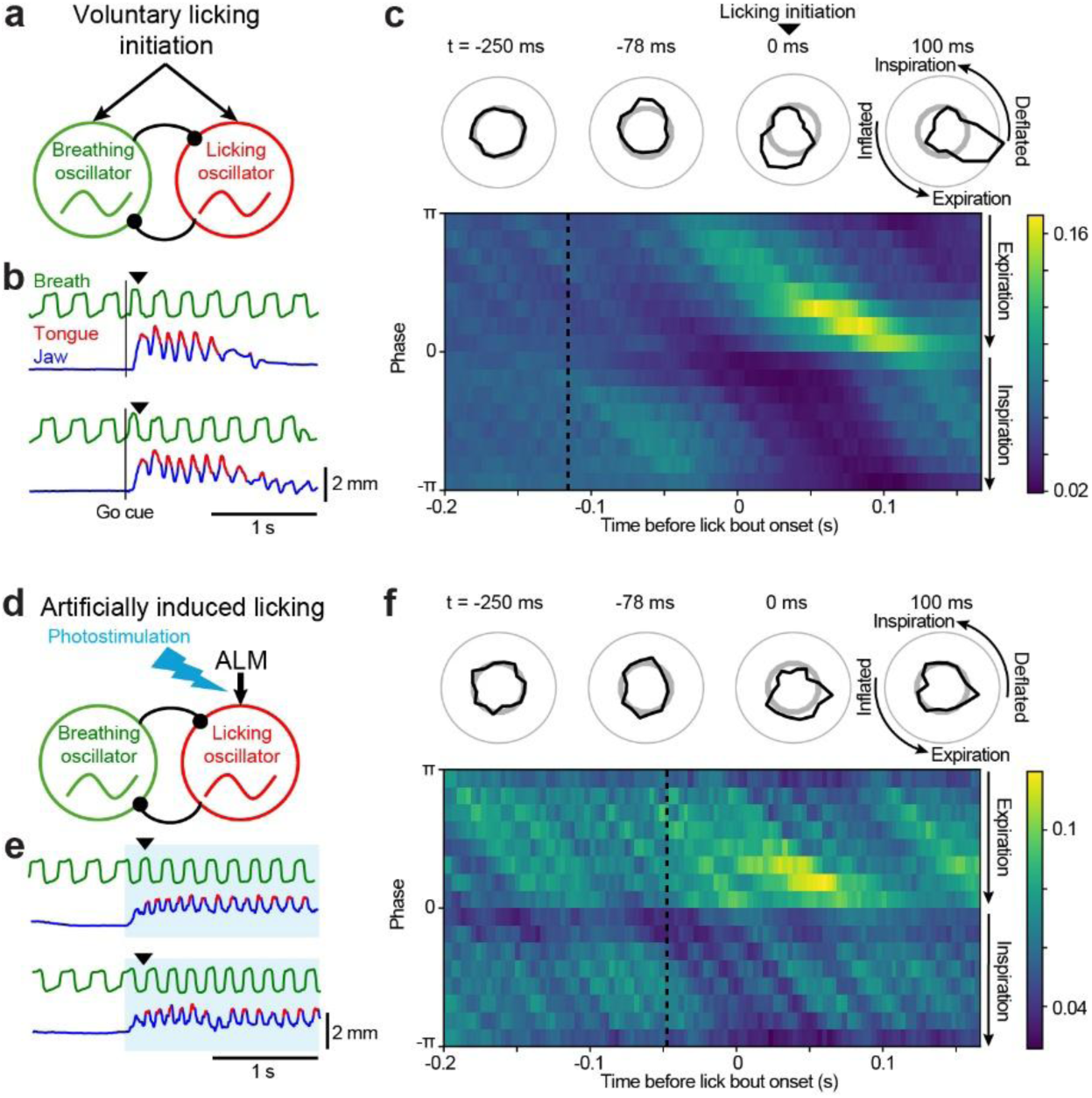
Coordination of licking initiation and breathing. **a**) Schematic of licking initiation that influences both breathing and licking oscillators. **b**) Example breathing (green), jaw movement (blue), and tongue protrusions (red) showing that breathing frequency is altered at licking initiation. Vertical line, Go cue onset. Arrow, onset of the first lick. **c**) Pre-emptive adjustment of breathing during licking initiation. *Top,* the polar plots show the distribution of breathing phase around the initiation of a licking bout. Bottom, the phase distributions plotted as a function of time relative to the onset of licking. Significant deviation from random distribution is detected at 115 ms before lick onset (dashed line), P<0.0001, two sample Kuiper test with Bonferroni correction. **d**) Artificially activating the licking oscillator by photostimulating ALM. **e**) Example breathing, jaw movement, and licking traces for artificially induced licking. Cyan, photostimulation. **f**) Breathing phase relative to licking onset for artificially induced licking. Same as **c**. Significant deviation of breathing phase is detected at 47.6 ms before lick onset (dashed line).

We tested whether this pre-emptive adjustment of breathing rhythm was recapitulated in artificially induced licking caused by ALM photostimulation (**Figure 5d**). Photostimulation evoked licking with a latency of 230.7 ± 31.5 ms (**Figure 5e**), which was significantly longer than voluntarily initiated licking (p<0.001, paired t-test, two-tailed). We aligned the breathing phase to the first tongue protrusion after photostimulaton (**Figure 5f**). Artificially induced licking did not recapitulate the pre-emptive breath adjustment observed before voluntary licking. Breathing phase was altered only shortly before licking onset (**Figure 5f**; 47.6 ± 10 ms), a significantly shorter latency than the pre-emptive adjustment before voluntary licking (**Figure 5g**; P<0.0001, bootstrap). These results uncovered additional coordination of licking and breathing beyond the autonomous cycle-by-cycle coordination by the brainstem circuits. ALM is necessary for licking initiation ^56^, but the 20 Hz artificial stimulation likely interfered with ALM’s coordination with top-down breathing pathways. These results suggest a role for these descending inputs in patterning the breathing oscillator at moments of licking initiation.

## Discussion

Our study outlines brainstem dynamics controlling structured breathing, licking, and swallowing during drinking behavior in mice (**Figure 1**). Using Neuropixels recordings, we map activity in the brainstem of behaving mice (**Figure 2**), which revealed distinct clusters of neurons activated during different orofacial rhythms with rich encoding of movement parameters. We further delineate a brainstem premotor network for licking with rhythm-generation properties (**Figure 3**). We found that coupled brainstem oscillators autonomously organize licking and breathing rhythms over rapid timescales (**Figure 4**). At moments of movement initiation, brainstem oscillators are further patterned (**Figure 5**), presumably by descending control.

Despite pharmacological, neuroanatomical, and manipulation studies, how brainstem circuits coordinate orofacial behaviors remains obscure, mainly because of extremely limited neurophysiological data. Electrophysiological studies have mostly focused on individual oscillators ^3,6,23^. In rodents, an oscillator controlling rhythmic whisking interacts with the breathing oscillator to coordinate breathing and whisking ^17,18^. But circuits for coordinating rhythmic ingestive behaviors remain unresolved. Here we dissect coupled brainstem oscillators coordinating breathing and drinking in behaving animals.

Brainstem circuits generate the rhythms underlying various orofacial behaviors, including suckling, chewing, and vocalization ^3,7,8^. These orofacial rhythms are also coordinated with breathing ^12,19,21^. For example, in humans breathing and speech are coordinated so that vocalization occurs only on exhalation ^57^, which is achieved by millisecond timescale coordination of respiratory and articular muscles ^58^. We find that mice temporally organize their licking and breathing cycle-by-cycle and this coordination is autonomously patterned by coupled brainstem oscillators (**Figure 4**). Rapid timescale coordination of other orofacial movements with breathing may be similarly governed by coupled brainstem oscillators.

Swallowing inhibits licking and breathing (**Figure 1k-n**), suggesting additional coupling of the swallowing pattern generator to the licking and breathing oscillators. Swallowing involves airway closures by the soft palate and epiglottis to avoid aspiration ^1,27^. Circuit coupling is necessary to provide protection from aspiration. Swallowing is thought to be controlled by a dorsal premotor network near NTS and a ventral premotor network near nucleus ambiguus ^6^ (**Figure 2h**). The circuits that connect these premotor networks to the licking and breathing oscillators remain to be elucidated ^59^. Discoordination of swallowing and breathing can cause choking, which is a leading cause of death among preschool children and elderly ^13^. There is a strong association of choking with numerous neurological conditions, including Alzheimer’s disease, dementia, Schizophrenia, Parkinson’s disease, and Rett Syndrome ^14–16^. Despite diverse causes, these symptoms may trace their roots to abnormal brainstem dynamics. Identification of these brainstem circuits could provide novel entryways for interventions.

The brainstem oscillators are further subject to descending control ^60^. Humans pre-emptively adjust breathing before vocalization ^57,61^. We find that mice pre-emptively adjust their breathing rhythm before licking. Our optogenetic manipulation experiment suggests contributions from sources beyond the brainstem for pre-emptive breath adjustment (**Figure 5**). Interestingly, ALM and latSC projections avoid the breathing oscillator (**Figure 3 and Extended Data Figure 9**). Descending control likely results from interactions of licking motor centers with top-down breathing pathways ^60^. Simultaneous recordings in these motor centers and brainstem oscillators may shed light on the descending signal that patterns brainstem oscillators.

A brainstem licking oscillator has been previously postulated ^5,28^. Licking could be evoked by stimulating *Phox2b^+^*neurons in the IRN ^9^. IRN is a large structure with diverse functions ^18,19,21,26,29–31^. Using photostimulation of *Phox2b^+^* neurons, we found a posterior-dorsal subregion of IRN and the adjacent subregion in PARN with autonomous rhythm-generating properties capable of driving sustained licking. It is unclear if the posterior-dorsal IRN/PARN generates rhythms intrinsically or through interactions with a broader network. The effect of stimulation could spread beyond the targeted neurons. Notably, the tongue-jaw premotor neurons and licking-related activity span a considerably broader region than the identified posterior-dorsal IRN/PARN region ^43,62^ (**Figure 3a-b**). Known descending licking pathways also target more extensive regions of the brainstem (**Figure 3c-f**) ^42,44–51,63^. The precise borders of the licking oscillator remain to be mapped with loss-of-function experiments. Moreover, the molecular profile of the licking oscillator neurons remains to be determined.

## Acknowledgements

We thank Rich Mooney, Hidehiko Inagaki, Kevin Franks, Yujin Han, and Jeff Yau for comments on this manuscript. We thank SWC Advanced Microscopy for providing technical assistance and maintenance of the serial block-face 2-photon microscope; Tim Harris Lab at Janelia Research Campus for support with the Neuropixels probe recordings and spike sorting. This work was supported by Simons Collaboration on the Global Brain, a Canadian Institutes of Health Research Postdoctoral Fellowship, a Taiwanese GSSA scholarship, the National Institutes of Health Grant NS142432, NS131229, NS132025, NS137920 and NS107466, Pew Scholar Program, the McKnight Foundation, the Howard Hughes Medical Institute, including from the Janelia Research Campus Visiting Scientists Program. Schematics in Figures 2 and 3 diagrams were created with BioRender.com.

## Author contributions

LDL, KS and NL conceived and designed the experiments. LDL and HK performed the behavioral experiment. LDL performed the electrophysiology experiments. HK performed the photostimulation experiments. AF contributed the directional licking task. SW contributed the imaging and brain alignment pipeline for electrode localization. RG contributed to the neurophysiology data analysis pipeline. AT and SLT contributed the ALM and superior colliculus anatomy data. SML and DK contributed the licking and breathing behavioral data in rats. LDL, HK, KS, and NL analyzed the data. HK, LDL, KS, and NL wrote the manuscript.

## Supplemental movies

**Supplemental movie 1.** Example video with tongue, jaw, and breathing tracking.

**Supplemental movie 2.** Example video showing jaw locking with no tongue protrusion evoked by a brainstem photostimulation site.

**Supplemental movie 3.** Example video showing jaw locking with transient tongue protrusions by a brainstem photostimulation site.

**Supplemental movie 4.** Example video showing rhythmic licking evoked by a brainstem photostimulation site.

## Methods

### Mice

This study is based on data from 72 mice (age >P60). 31 VGAT-ChR2-EYFP animals (22 male, 9 female; JAX 014548, >P60) were used for electrophysiology in the brainstem. 9 Sim1_KJ18-Cre (MMRRC 031742) and 2 Tlx_PL56-Cre (MMRRC 036547) crossed to Ai32 (Rosa26-ChR2 reporter mice, 3 male, 8 female, JAX 012569) were used for ALM photostimulation. 1 wildtype mice with pAAV-hSyn-hChR2(H134R)-EYFP injected in the lateral superior colliculus (SC) and 2 Vglut2-ires-Cre mice with a Cre-dependent ChR2 virus injected in the lateral SC were tested for latSC photostimulation to evoke licking. 11 Phox2b-Cre mice (JAX 016223) and 16 Phox2b-Cre crossed to Ai32 mice were used for brainstem photostimulation. 7 Long-Evans adult female rats (Charles River Laboratory) with weights ranging from 300 to 424 grams were used for behavioral studies.

All animal procedures were approved by the Institution Animal Care and Use Committee at Baylor College of Medicine, Duke University, and the University of California, San Diego. Mice were housed in a 12:12 reversed light/dark cycle and tested during the dark phase. On days not tested, mice received 0.5–1 ml of water. On other days, mice were tested in experimental sessions lasting 1–2 h where they received all their water (0.5–1 ml). If mice did not maintain a stable body weight, they received supplementary water ^64^. All mice surgical procedures were carried out aseptically under 1–2% isoflurane anesthesia. Buprenorphine Sustained Release (1 mg/kg) and Meloxicam Sustained Release (4 mg/kg) were used for preoperative and postoperative analgesia. A mixture of bupivacaine and lidocaine was administered topically before scalp removal. After surgery, mice were allowed to recover for at least 3 days with free access to water before water restriction. Surgical procedures on rats were carried out under anesthesia induced by ketamine (40-100 mg/kg) and xylazine (5-10 mg/kg) injection. Supplemental administration of ketamine and xylazine was administered if needed and buprenorphine (0.03 - 0.05 mg/kg) was administered for post-op analgesia. Post-op rats were allowed to recover for at least two days.

### Surgery

The details of the headbar implantation surgery in mice have been previously described (dx.doi.org/10.17504/protocols.io.bcrsiv6e). Briefly, the skin and periosteum above the skull were cut away, and a layer of cyanoacrylate (Krazy glue) was used to adhere the headbar to the skull.

We injected Cre-dependent ChR2 or ChRimine viruses into the brainstem of Phox2b-Cre mice for photostimulation experiment to map the licking oscillator. 250 nL of AAV8.Ef1a.DIO.ChRmine.mScarlet.Kv2.1.WPRE (Stanford Viral Core, GVVC-AAV-188, 8.44e12 GC/ml), AAV1.CAGGS.Flex.ChR2-tdTomato.WPRE.SV40 (UPenn Viral Core, titer, 1.38e13 GC/ml), or AAV9.EF1a.double floxed.hChR2(H134R).mCherry.WPRE.HGHpA (Addgene, 20297, 1e13 GC/mL) was injected into the brainstem, followed by implantation of a 5-mm long optical fiber (Thorlabs, CFML12L05) over the injection site. The injection coordinate was posterior 0.7-3.2 mm from lambda, lateral 0.5-1.9 mm, depth 4.2-5 mm. The injection was made through the thinned skull using a piston based volumetric injection system. Glass pipettes (Drummond) were pulled and beveled to a sharp tip (outer diameter of 30 mm). Pipettes were back-filled with mineral oil and front-loaded with viral suspension immediately before injection. In Phox2b-Cre x Ai32 mice, an optical fiber was implanted across a similar range of stereotaxic coordinates (posterior 1.7-3 mm from lambda, lateral 0.5-1.3 mm, depth 3.9-5 mm) to target the brainstem for photostimulation. In a subset of mice, separate virus injection and fiber implant was performed on each hemisphere to test two different brainstem locations.

For latSC photostimulation experiments, we injected 300 nL of AAV1.CAGGS.Flex.ChR2-tdTomato.WPRE.SV40 (UPenn Viral Core, titer, 1.38e13 GC/ml) in Vglut2-ires-cre mice or AAV2. hSyn.hChR2(H134R).EYFP in wildtype mice. We targeted a lateral region of SC associated with licking (posterior 3.5 mm from bregma, lateral 1.5 mm, depth 2.5 mm) in either the left or right hemisphere ^48,49,51^.

Chronic surgical procedures on rats for measuring breathing were described in detail in prior work ^65^. In brief, an incision was made along the midline on the skull and the nasal bone. Soft tissues were removed from the exposed surface. Head screws (#00-90, McMaster-Carr) were implanted onto the skull and a hole, 1 mm in diameter, was drilled 2 mm from the rostral end of the nasal bone and 2 mm from the midline. A thermocouple (5TC-TT-K-36-36, Omega) was inserted into the hole, position away from the walls, with the wires attached to the skull and nasal bone with dental cement. A lightweight head holder (0.85” x 0.5”) was attached to the skull with dental cement.

### Histology

Mice were perfused transcardially with PBS followed by 4%paraformaldehyde (PFA)/0.1MPBS. The brains were fixed overnight and transferred to 20% sucrose before sectioning on a vibratome (Leica). Coronal 50 mm free-floating sections covering the hindbrain were collected. Slide-mounted sections were imaged on an Olympus Macroscope.

We identified the coronal section where the optical fiber tips are located and aligned the coronal section to the Allen Mouse Common Coordinate Framework (CCF) using landmark-based image registration ^66^. The registration target was the 10 μm per voxel CCF anatomical template. To align a coronal section, we first manually selected the coronal plane in the anatomical template that best corresponded to the section. Next, we manually placed control points at corresponding local landmarks in each image (**Extended Data Figure 9e-f**). The image was warped to the CCF using an affine transformation followed by a non-rigid transformation using b-splines ^67^. Images were warped using the B-spline Grid, Image and Point based Registration package available on the Matlab FileExchange (https://www.mathworks.com/matlabcentral/fileexchange/20057-b-spline-grid-image-and-point-based-registration). We performed this procedure independently for each brain section.

### Multi-directional licking task

Electrophysiological recordings from the brainstem in mice were performed in a directional licking task. Mice were conditioned on the multi-directional licking tasks with 9 targets (**Figure 1a**). The horizontal distance between the adjacent targets was 3.5 mm and the vertical distance was 1.5 mm. Each trial began with a drop of water (2 µL) dispensed from the lick spout and the lick spout immediately moved to one of the nine targets. The mice had 4 seconds to lick the water from the lick spout before it was retracted to a position unreachable by the mice. The mice performed 40 trials at each target position. Each trial was at a pseudo-random position with 2 seconds of intertrial interval. The mice were conditioned for 1-3 days until they accurately reached all 9 targets within 4 seconds, and their licks were reliably detected by the piezoelectric lick spout (<10% of trials without any licks detected). Electrophysiological recordings only commence after the mice could accurately lick to the targets with less than 10% miss rate.

### Cued licking task

ALM and brainstem photostimulation experiments in mice were performed in a cued licking task. A lickspout was placed at a fixed location in front of the mice to deliver water rewards and record licks. At the beginning of each trial, an auditory Go cue was given (pure tone, 3.4 kHz, 0.1 s duration), after which mice could lick the lickspout to trigger a water reward. Trials in which mice did not lick within a 0.5 s window after the Go cue were rare and were counted as misses. The inter-trial-interval was random (4.5 - 6s). After the trial start, the Go cue was presented after a random wait time between 0.5 and 2s, making the Go cue timing unpredictable. To prevent impulsive licking, the wait time counter restarted if mice licked early, within a 0.5 s window prior to the Go cue. On a random subset of trials (25-50%), the Go cue was substituted with photostimulation. Water reward was withheld on photostimulation trials.

### Photostimulation

For ALM photostimulation in mice, light from a 473-nm laser (UltraLasers, MBL-FN-473-300mW) was controlled by an acousto-optical modulator (AOM; Quanta Tech, MTS110-A3-VIS), and focused onto the skull or brain surface (beam diameter: 400 µm at 4σ). The photostimulus was given at 20Hz (duration: 2 s; average power: 1.6 mW, 20% duty cycle).

For brainstem photostimulation in mice, light was delivered to the brainstem through an optical fiber (Thorlabs, 200µm core, 0.39NA, Part No. CFMLC12L05) coupled to either a 473-nm laser for ChR2 stimulation (UltraLasers, MBL-FN-473-300mW) or a 635 nm laser for ChRmine stimulation (MRL-III-633–50, Ultralaser). The photostimulus was constant (duration: 2 s; average power: 5 mW).

For latSC photostimulation in mice, light was delivered to latSC through an optical fiber (Thorlabs, 200µm core, 0.39NA, Part No. CFMLC12L02) coupled to a 473-nm laser (UltraLasers, MBL-FN-473-300mW). The photostimulus was a 40 Hz sinusoid (duration: 1.3 s; average power, 8 mW). Mice were tested in absence of any task. The mice were also not water restricted.

### Videography and tracking of breathing in mice

Breathing was tracked by placing an airflow meter (Honeywell AMW3300V) in front of the mouse’s nose (**Figure 1b**). The change in air pressure is converted to voltage simultaneously recorded with electrophysiology and videos at 25kHz in SpikeGLX. Orofacial movements were tracked at intervals of 3.4 ms by CMOS cameras (Blackfly, FLIR) from the side and bottom views (**Figure 1b**). Each frame of videos is triggered and acquired at 3.4 ms intervals with custom software (https://github.com/LiuDaveLiu/pySpinCapture/tree/master). The bottom view was acquired at 720 pixels X 540 pixels, while the side view was acquired at 400 pixels X 480 pixels. The videos were acquired in the dark with the mouse illuminated from multiple directions with infrared lamps (940 nm). The 2 views were calibrated to track orofacial features in 3D using a Matlab toolbox (https://github.com/nghiaho12/camera_calibration_toolbox_octave).

### Behavioral data analysis in mice

Breathing phase was extracted from Hilbert transform of the airflow measurement. The onset of inspiration was defined as the timepoint where the phase was equal to 0.

The jaw and tongue movements were tracked by DeepLabCut (DLC, https://github.com/DeepLabCut). 5000 frames from the bottom view and 5000 frames from the side view were manually labeled for the tip of the jaw, tip of the tongue, and the lickport (**Figure 1b**). The frames were used to train separate networks for the bottom and side views. The tracking of jaw, tongue and lickport were transformed to trajectories in 3D using a rigid body transform in the toolbox above. The onset of lick was defined as the first video frame in which the tongue was detected. Lick angle was defined as the angle between the tip of the tongue at the fully protracted state to the resting position of the jaw (**Extended Data Figure 8g**).

Swallowing events were identified from breathing traces using a heuristic algorithm that detects breathing pauses during expiration (**Extended Data Figure 1a**). First, the algorithm identified the negative inflection points in breathing traces using the Python function *find_peaks()*. Next, only the peaks during the expiration phase of breathing were kept. Because swallowing occurs during licking, the detected events were further restricted to occur within a lick bout. Finally, the breathing pauses containing the detected events needed to exceed 50 ms in duration (defined as the period in which breathing trace velocity was below a threshold). The swallowing onset time was defined as the first time point of the breathing pause. Although swallowing was identified from breathing, we found licking was also inhibited by swallowing. As a control, we also tested if the alignment of licking and swallowing could accidentally arise from the phase-coupling of licking and breath (**Extended Data Figure 1b-c**).

### Simulation of licking-breathing coupling

We simulated the expected lick and breathing timings for a free-running breathing oscillator that unidirectionally adjusts the timing of licking (**Extended Data Figure 1d-f**). First, inspiration timing was generated by drawing inter-breath-intervals from a distribution of breathing frequency based on empirical data in mice (**Figure 1d**). Next, lick timing was set by placing each lick at a fixed latency before each inspiration. This model was based on the empirical observation that licking occurred at a fixed latency before each inspiration (**Figure 1f**). Next, for inter-lick-intervals exceeding the empirically observed range, additional licks are added in-between the existing licks to maintain a licking frequency commensurate with the data (**Figure 1d**). Gaussian noise (mean, 0; std, 10ms) was added to the simulated lick timing. Finally, we analyzed lick timing and breath timing from the simulation relative to each other.

We also simulated the expected lick and breathing timings for a free-running licking oscillator that unidirectionally adjusts the timing of inspirations (**Extended Data Figure 1g-i**). First, lick timing was generated by drawing inter-lick-intervals from a distribution of licking frequency based on empirical data (**Figure 1d**). Next, inspiration timing was set by placing inspirations at a fixed latency before each lick. Additional constraint was applied such that inter-breath interval must fall within the empirically observed range (**Figure 1d**) and inspiration was omitted otherwise. This model was based on the observation that inspiration timing was influenced by the timing of upcoming licking (**Figure 1h**). Gaussian noise (mean, 0; std, 10ms) was added to the simulated inspiration timing.

### Licking and breathing in freely moving rats

To record licking in freely moving rats, water-restricted rats were placed into an acrylic box where they could access a water bottle and lick for water. Contacts of the tongue to the water bottle spout were detected by the lickometer (designed by ^68^). The implanted thermocouple was connected to a preamplifier (DAM80, World Precision Instruments). The output signals from the thermocouple and the lickometer were sampled by the data acquisition system (PowerLab, ADInstruments) at 40 kHz using the software LabChart (ADInstruments). The session ended when the animal no longer showed the intent to lick from the water bottle.

### Licking and breathing data analysis in freely moving rats

Licking data with noisy breathing signals and additional ∼188 licks (<0.5% of total 39k licks used in analysis) from short bouts between 0.35 and 1.75 s were excluded from analysis. The thermocouple signal was first low pass filtered at 50 Hz (Butterworth, fifth order) and down sampled to 2.0 kHz along with the lickometer signal for downstream analysis. From the licking signal, events of contact (or release) by the tongue to (or from) the waterspout were identified by the rising and falling edges in the bi-level signal. For plotting the raster (volcano) plots (**Extended Data Figure 2**), the breathing signal is high pass filtered at 1 Hz (Butterworth, third order) and low pass filtered at 20 Hz (Butterworth, third order), followed by the extraction of the inspiration onsets by methods adopted from ^65^ with a threshold of a rise to 10 % of onset amplitude. Data analysis was done using MATLAB (MathWorks).

### Electrophysiological recording with Neuropixels probes

Detailed procedure for electrophysiological recording can be found elsewhere ^36^. Briefly, after mice were conditioned on the multi-directional licking task. We made craniotomies of 1 mm diameter on both left and right sides of the brain. In each daily recording session, 1-2 Neuropixels 1.0 probes were acutely lowered through the craniotomies to the target regions at 5 µm/s with micromanipulators (Sensapex). Prior to the probe insertion, the probe tips were painted with CM-DiI (Thermo Fisher) to track their location in the brain (dx.doi.org<u>/10.17504/protocols.io.wxqffmw</u>). While the mouse performed the task, recordings were collected from the 384 electrodes closest to the tip of the probe, i.e. bank 0. At the end of each session, we recorded from the adjacent 384 electrodes, i.e. bank 1, to estimate the surface of the brain (**Extended Data Figure 3c**). Online, the electrophysiological recordings were high-pass filtered at 300 Hz and sampled at 30 kHz using SpikeGLX (https://billkarsh.github.io/SpikeGLX/). Daily recording sessions lasted 1-2 hours. At the end of each recording session, we retracted the probe out of the brain, and the craniotomy is sealed with removable adhesive (Kwik-Cast, World Precision Instruments) and opened again prior to the next session of recording. Typically, we performed 4-5 probe insertions per craniotomy across days. The insertions were spaced at least 250 µm apart to clearly separate insertions across sessions. Neuropixels probes span the entire dorsal-ventral range of the medulla and pons (**Figure 2a-c**). Across multiple recordings from multiple mice, we varied craniotomy locations along medial-lateral and anterior-posterior axes to sample from the entire medulla and a large portion of pons (**Figure 2d**).

### Preprocessing, spike sorting, and quality metrics

Raw signals are high-pass filtered at 300 Hz using a fourth-order Butterworth filter in both forward and backward directions. We performed spike sorted in Kilosort 2.5 (https://github.com/MouseLand/Kilosort/releases/tag/v2.5.2). We used a set of quality metrics (https://doi.org/10.25378/janelia.24066108.v1) to filter out units for further analyses ^35^: Amplitude > 150 µV, ISI violation < 10, spike rate > 0.2 Hz, presence ratio > 0.9, amplitude cutoff < 0.15 and drift metric < 0.5. The workflow for preprocessing, spike sorting, and quality control metrics can be found here (https://github.com/jenniferColonell/ecephys_spike_sorting).

After applying these quality control metrics to filter sorted units (60 ± 22 units for each Neuropixel penetration, mean ± std across recordings), we obtained 20,300 clear single units, including 5,700 units in the cerebellum, 3,700 units in the pons, and 10,900 units in the medulla. Spike rate of units differed widely across brain regions. Neurons in the cerebellum and the vestibular nucleus exhibited higher spike rates than neurons in the medulla (22 ± 29 spikes/s, mean ± std, for cerebellum, 38 ± 32 spikes/s for vestibular nucleus, and 11 ± 14 spikes/s for medulla, **Figure 2e**). To rule out possible movement artifacts during licking, we examined the unit activity during licking and confirmed that it was unaffected by the quality of spike sorting (**Extended Data Figure 4**).

### Electrode localization

The electrode localization workflow is described in detail elsewhere ^36^. After the last recording session for each mouse, we transcardiacally perfused the mouse with saline followed by 4% paraformaldehyde and dissected out the brain for imaging. The brains were postfixed in 4% paraformaldehyde at room temperature for 3 days before transferring to PBS at 4°C for 3 days. The postfixed brains were embedded in agarose before imaging. The brains were imaged with the custom serial block-face 2 photon microscope (SBF2P) at the Sainsbury Wellcome Centre ^69,70^. The brains were section at 50 µm interval and illuminated with a Chameleon Ultra I two-photon laser at 920 nm (110 mW). We acquired images 2 depths of 25 µm per section in two channels. The brains autoflurescence was captured at 425-495 nm, and the fluorescence from DiI at 570+ nm. The microscope was controlled by (ScanImage v5.6, Vidrio Technologies, USA) using BakingTray, a custom software wrapper for setting up the imaging parameters (https://github.com/SainsburyWellcomeCentre/BakingTray, https://doi.org/10.5281/zenodo.3631609). Images were assembled using StitchIt (https://github.com/SainsburyWellcomeCentre/StitchIt, https://zenodo.org/badge/latestdoi/57851444).

The images collected with the SBF2P microscope were registered to the Allen anatomical template in CCF v3 (https://github.com/int-brain-lab/brainregister) via an Elastix-based software (Klein et al., 2010). The Elastix parameters used were similar to that from previous studies ^36,70^, the affine transform parameters was optimized at four resolutions and the B-spline transform parameters optimized at six resolutions ^71^. The optimization metric is based on the Advance Mattes mutual information that minimizes the discrepancy between the moving and the fixed images ^72^. The adaptive stochastic gradient descent was done iteratively at a maximum of 500 iterations at each resolution.

After registering the brain images to the CCF space, we manually annotated along the probe tracks labelled by DiI fluorescence to recover the probe tracks in CCF (**Figure 2d and Extended Data Figure 3b**). The electrodes are assigned to the probe track by anchoring electrodes to electrophysiological landmarks that mark the transition between brain regions. Specifically, the electrophysiological landmarks used are surface of the cerebellum, the transition between the fourth ventricle and the medulla, and the transition between the vestibular nucleus and other parts of the medulla (**Extended Data Figure 3c**). There is usually a noticeable difference in the spike count in these brain transitions (**Figure 2e and Extended Data Figure 3d**). After anchoring the electrodes on the probe corresponding to the electrophysiological landmarks, the remaining electrodes were linearly interpolated or extrapolated by calculating a scaling factor to scale the interelectrode distance between landmarks ^36^. Since units recorded on the probe can be assigned to an electrode where they have the highest waveform amplitude, localizing electrodes is the equivalent of localizing units in the CCF.

### Electrophysiology data analysis

To quantify neuronal selectivity for licking, breathing, or swallowing, we calculated a modulation index for each unit. For licking and breathing, modulation index was the spike rate difference between a neurons’ preferred phase and non-preferred phase. To obtain phase tuning, the jaw position was bandpass filtered between 3 and 15 Hz with a 4-pole Butterworth filter run in forwards and backwards direction, and breathing was bandpass filtered between 1 and 10 Hz ^18,73^. Phase was extracted from Hilbert transform of the filtered jaw movement traces and the breathing traces (**Extended Data Figure 5a**). The instantaneous phase of the jaw and breathing where spikes occurred were binned and normalized to get a phase tuning curve for each unit. The tuning, *r*(*x*_*p*_), was fitted to a circular Gaussian using a non-linear least squares algorithm (optimize.curve_fit in SciPy),

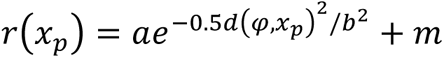

Where *a* scales for the amplitude of the tuning curve, *φ* is the preferred phase of unit (**Extended Data Figure 8b** and **e**), *x*_*p*_ is the phase of the jaw movement or breathing, and *d*(*φ*, *x*_*p*_) is the shortest circular distance between *φ* and *x*_*p*_. *b* scale for the width of the tuning curve and *m* is the baseline. Licking and breathing modulation index (MI) was defined as *MI* = (*r*(*φ*) − *r*(*φ* − *π*))/*r*(*φ*). For each unit, we quantified the significance of the MI by randomly sampling 1000 times from the jaw movement or breathing phases for that session. This resampling generates a distribution of 1000 MIs for each unit to compare the measured MI against. High modulation index units were defined as P < 0.001 from the Wilcoxon rank sum test against resampled distribution and MI >0.6. The number of licking and breathing modulated units varied as a function of the MI criterion, but their relative proportion was robust to the criterion.

For swallowing, the MI was the spike rate difference between the time window around swallowing (100 ms centered on swallow onset) and a 200-ms time window either before (-300 to -100 ms from swallowing onset) or after swallowing (100 to 300 ms), normalized to the peak spike rate during swallowing. MIs was calculated separately relative to the window either before or after swallowing, then averaged to obtain a single MI for each neuron. For each unit, we quantified the significance of the MIs using non-parametric bootstrap. In each round of bootstrap, we randomly sampled with replacement the single trial spiking data and recalculated the MI on the re-sampled data. Performing this procedure 1000 times generated a distribution of MIs. The p value for was the fraction of times the MI changed sign, e.g. if a unit had a positive MI, p value for this unit was the number of times resampling produced a negative MI.

To quantify the direction tuning of licking related units (**Extended Data Figure 8**), direction tuning curves were calculated by binning the response for each lick according to the lick angle of the lick.

### Generalized Linear Model (GLM)

To look at each tracked features’ contribution to the units’ spiking, we devised a GLM to fit to the response of each unit at 17 ms resolution (api.GLM in statsmodel). The exogenous variables are the tongue and jaw tracking in x, y, and z dimensions plus the breathing tracking. The exogenous variables are all median filtered to have temporal resolutions of 17 ms. The exogeneous variables are all z-scored to have equal mean and variance. For the tongue variable, periods where the tongue is not in the videos, as marked by the likelihood of the DLC tracking to be less than 0.05 in both views of the video, are set to 0. The endogenous variable for the GLM is the spike rate of each unit binned at 17 ms (**Extended Data Figure 7a**). The GLM used logistic link function with Poisson spiking statistics. We used every 5th trial in a session to make up the validation set and the remaining trials to be the training set. The GLM is fitted with the spike rate of each unit shifted at 5 lags of 17 ms in each direction. We used L1 regularization for the weights. We fitted the GLM parameters with the training set and used the parameters for each unit to predict the spiking rate of the corresponding validation set. The variance account for (VAF) of each unit at each lag is calculated as,

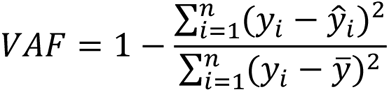

Where *y*_*i*_ is the trial concatenated spike rate at time bin *i*, and *ŷ*_*i*_ is the predicted spike rate. *n* is the total number of time bins in the session, and *ȳ* is the mean spike rate of the unit.

### Convolutional autoencoder (CAE)

To capture features of orofacial movements not captured with DLC tracking. We used a convolutional autoencoder (CAE) to represent the videos in a 32-dimensional embedding space (**Extended Data Figure 7e**). The architecture of the CAE has been previously described ^40^. The anto-encoder is first composed of two residual blocks each with three convolutional layers. The residual blocks are followed by two fully connected linear readout layers with output size 128 and 32. We trained one auto-encoder for each animal. The training set for each anto-encoder consists of 80% of trials from all sessions of that animal. Each frame of the videos is downsampled to a size of 120 pixels X 112 pixels for the input layer. Each CAE is trained for 20,000 iterations. We used L2 regularization to prevent overfitting. The GLM fitting procedure was similar to above but we replaced the DLC tracking features with the embeddings from the auto-encoder (**Extended Data Figure 7f**). The GLM is fitted with 80% of the data and cross-validated with 20% of the data.

In general, the variance accounted for by the embeddings of the CAE is higher than that from DLC tracking (**Extended Data Figure 7g**). For units in the preBötzinger complex, activity prediction using CAE led to slightly worse predictions than tracked features (**Extended Data Figure 7h**). The lower prediction could be due to occlusion of breathing-related movements of the nose by the nose cone in the video. This suggests a 1-dimensional signal as measured by airflow was sufficient to capture the activity of the breathing oscillator and including additional orofacial features from the rest of the face and body did not improve prediction power.

## Code availability

Code used for analyses will be deposited upon publication.

## Data availability

Data will be deposited upon publication.

