## Extended Data Figures for "A brainstem map of orofacial rhythms"

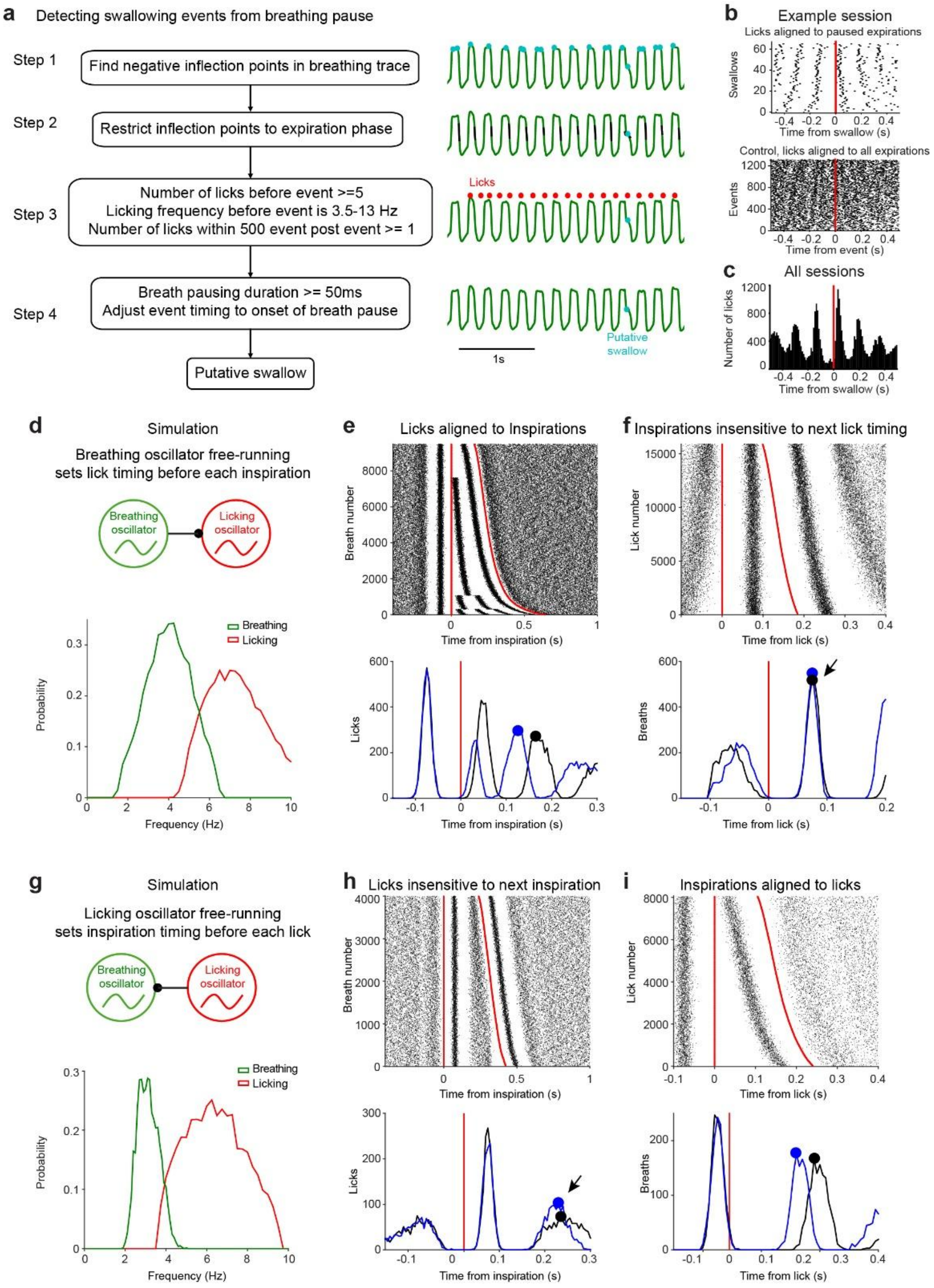


**Extended Data Figure 1. Detection of swallowing from breathing traces and lick-breath temporal relationship.**

- **a.** Algorithm to detect swallowing events from paused expirations.
- **b.** Example session. *Top,* lick timing aligned to the detected swallow events. The timing of each lick is represented by a dot plotted against swallowing onset (red lines). The licks are ranked by the inter-lick interval where the faster licking frequency is at the top. Licking was paused during swallowing. *Bottom,* lick timing aligned to all expirations as a control. Lick was not paused during expiration.
- **c.** Lick probably aligned to swallowing across all sessions.

- **d.** Simulation of a free-running breathing oscillator that sets the timing of licking (Methods). *Bottom*, histograms of licking and breathing frequencies from the simulation.
- **e.** Licks aligned to breathing. Compare to Fig 1f.
- **f.** Breaths aligned to licking. Compare to Fig 1h. Arrow points to a discrepancy with empirical data.
- **g-i.** Simulation of a free-running licking oscillator that sets the timing of inspirations. Same as **d-f**.

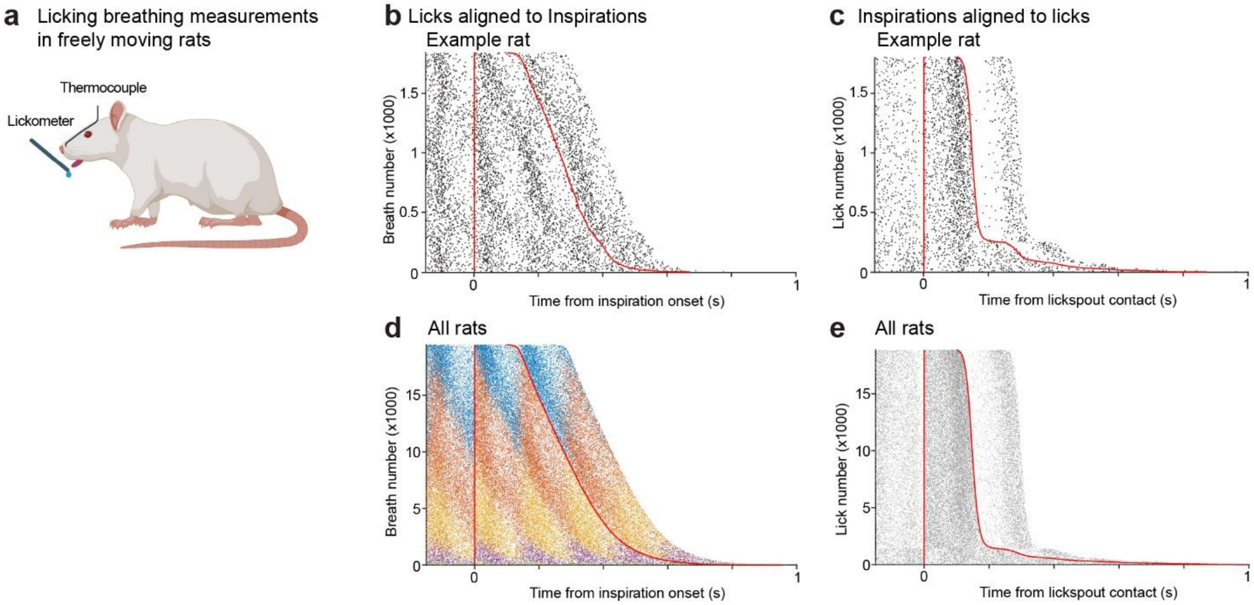


**Extended Data Figure 2. Lick-breath temporal relationship measured in freely moving rats.**

- **a.** Measurements of licking and breathing in freely moving rats. Licking is measured by contacts with lickometer. Respiration is recorded by an implanted thermocouple in the nasal cavity.
- **b.** Licks aligned to breaths. The timing of each lick is represented by a dot plotted against inspiration onset, red lines. The inspirations are ranked by the inter-breath interval where the shortest breath is at the top. Data from an example rat.
- **c.** Inspirations aligned to licks; same as panel **b**.
- **d.** Licks aligned to breaths. Different colors correspond to different modes of oscillation with 1 vs. 2 vs. 3 vs. 4 licks between breaths. Data from 7 rats.
- **e.** Inspirations aligned to licks. Data from 7 rats.

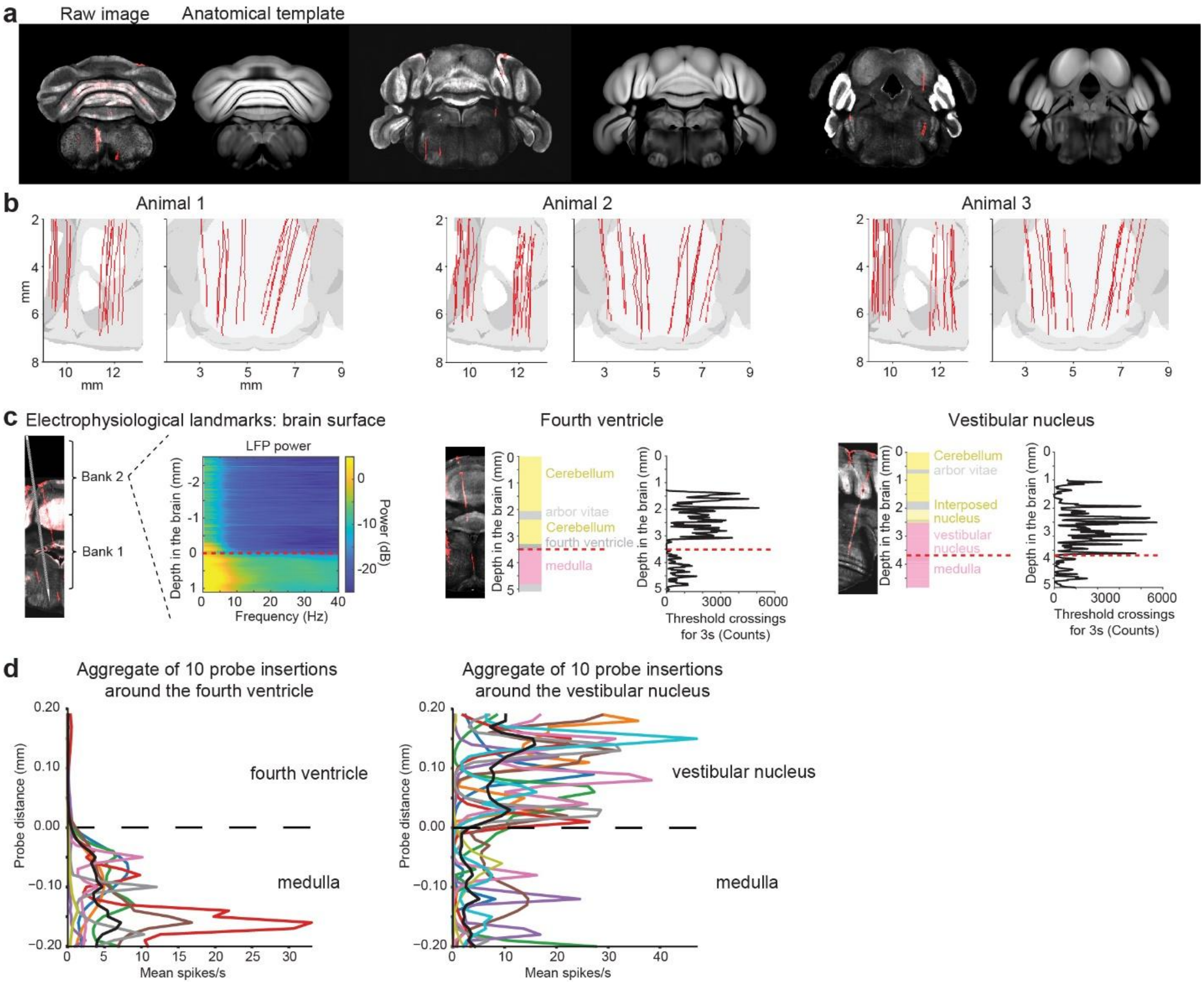


**Extended Data Figure 3. Examples from key steps of the electrode localization workflow.**

- **a.** Left, example sections from the serial block-face 2-photon microscope aligned to the Allen anatomical template. Right, the corresponding section of the anatomical template used for alignment.
- **b.** Three example brains with aggregate of probe tracks labeled in red. The outline of the Allen anatomical template is shaded in gray.
- **c.** Example electrophysiological landmarks used in electrode localization: the surface of the brain, the transition between the fourth ventricle and the medulla, and the transition between the vestibular nucleus and the medulla.
- **d.** Aggregate of mean spike rate for 10 probe insertions around the transition between the fourth ventricle and the medulla, and the transition between the vestibular nucleus and the medulla.

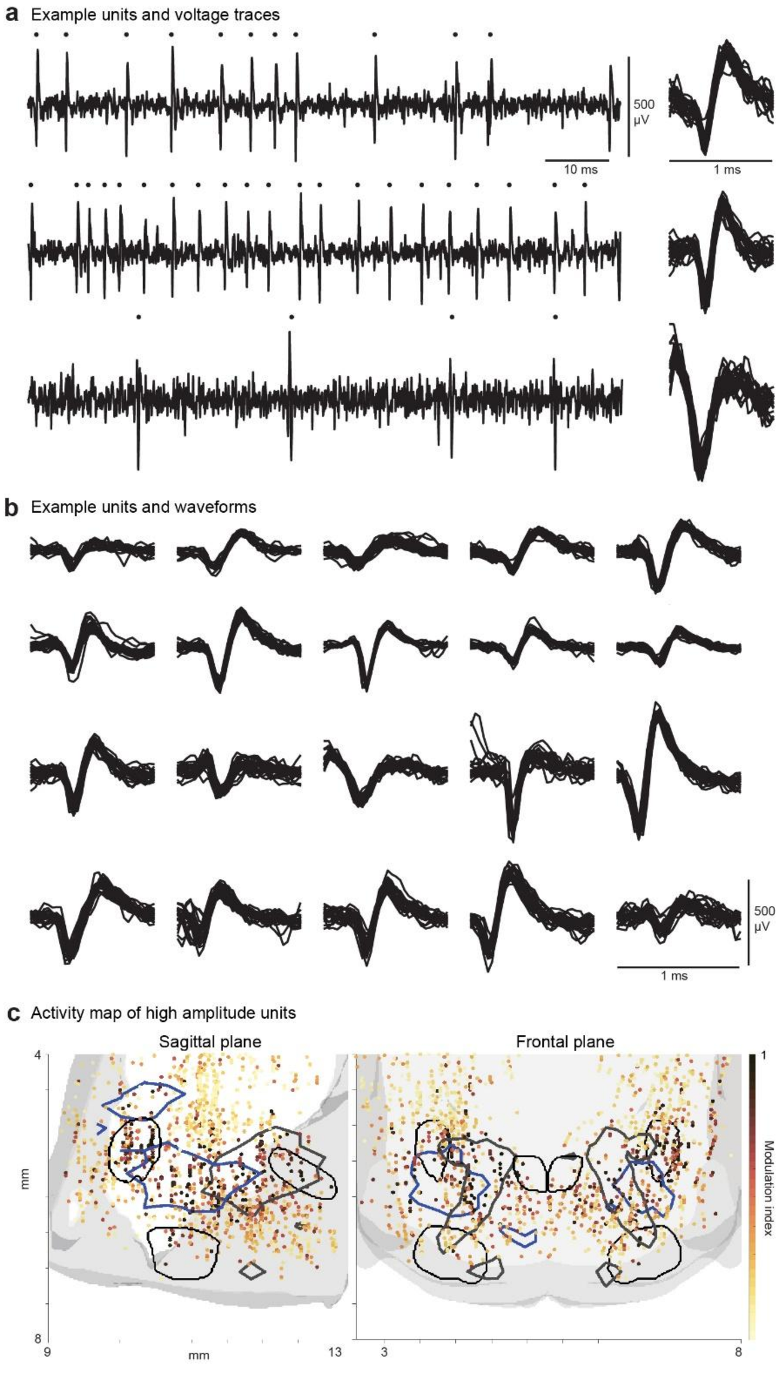


**Extended Data Figure 4. Example units after spike sorting.**

- **a.** Example single units from raw voltage traces. The voltage recordings are plotted on the left, and the spike times are indicated by dots on top. The corresponding waveforms are plotted on the right.
- **b.** Additional example units, each with 25 example waveforms.
- **c.** Licking activity map was insensitive to the quality of spike sorting. To obtain units with the highest amplitude, the waveform amplitude quality metric was increased from 150 µV to 500 µV. This filtered the 19,000 units with amplitude > 150 µV to 2223 unit with amplitude > 500 µV. The modulation index for licking is indicated by the hotness of the color. The location of the units is plotted against the premotor and motor nuclei for jaw and tongue muscles.

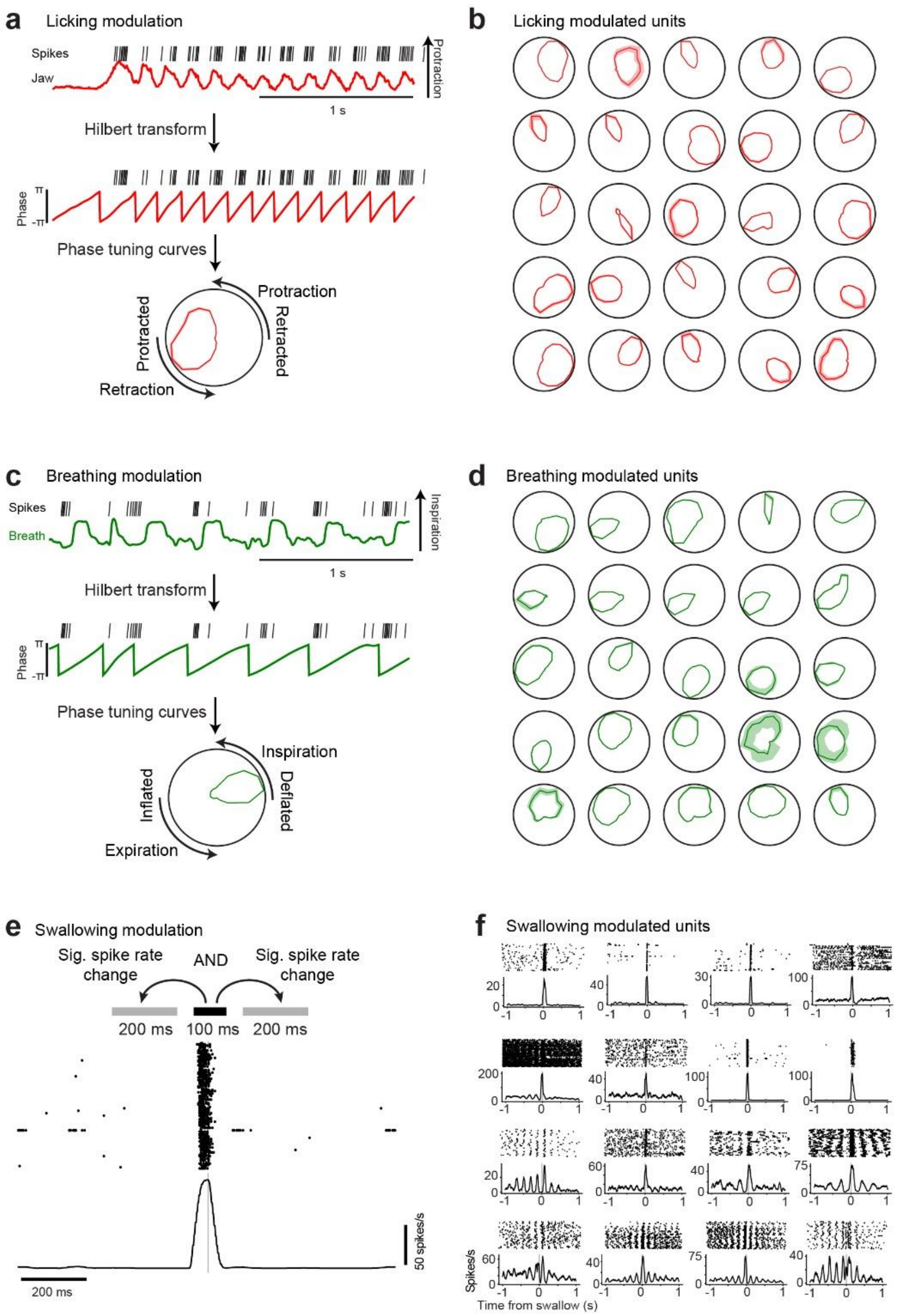


**Extended Data Figure 5. Tuning curves of example brainstem units.**

- **a.** *Top*, example jaw tracking with spikes as ticks on top. *Middle*, Hilbert transform to obtain phase from the licking traces. *Bottom*, the instantaneous phase where the spikes occurred are binned to create a tuning curve for each unit. Each tuning curve is normalized by the distribution of all phases in the session. Shadings are 95% confidence interval from resampling.
- **b.** Example licking tuning curves for 25 units. Shadings are 95% confidence interval from resampling.
- **c.** *Top*, example breathing tracking with spikes as ticks on top. *Middle*, Hilbert transform to obtain phase from the breathing traces. *Bottom*, the instantaneous phase where the spikes occurred are binned to create a tuning curve for each unit. Each tuning curve is normalized by the distribution of all phases in the session. Shadings are 95% confidence interval from resampling.
- **d.** Example breathing tuning curves for 25 separate units. Shadings are 95% confidence interval from resampling.

**e.** Raster and PSTH of an example units with swallowing related activity. Swallowing modulation was the spike rate difference between a 100 ms window centered on swallowing onset (gray line) relative to 200 ms windows before (-300 to -100 ms from swallowing onset) and after (100 to 300 ms). Significant swallowing modulation was calculated using bootstrap.

**f.** Example units with significant modulation during swallowing. *Top*, eight units show transient activity during swallowing. *Bottom*, eight units show licking-related activity which are further modulated during swallowing.

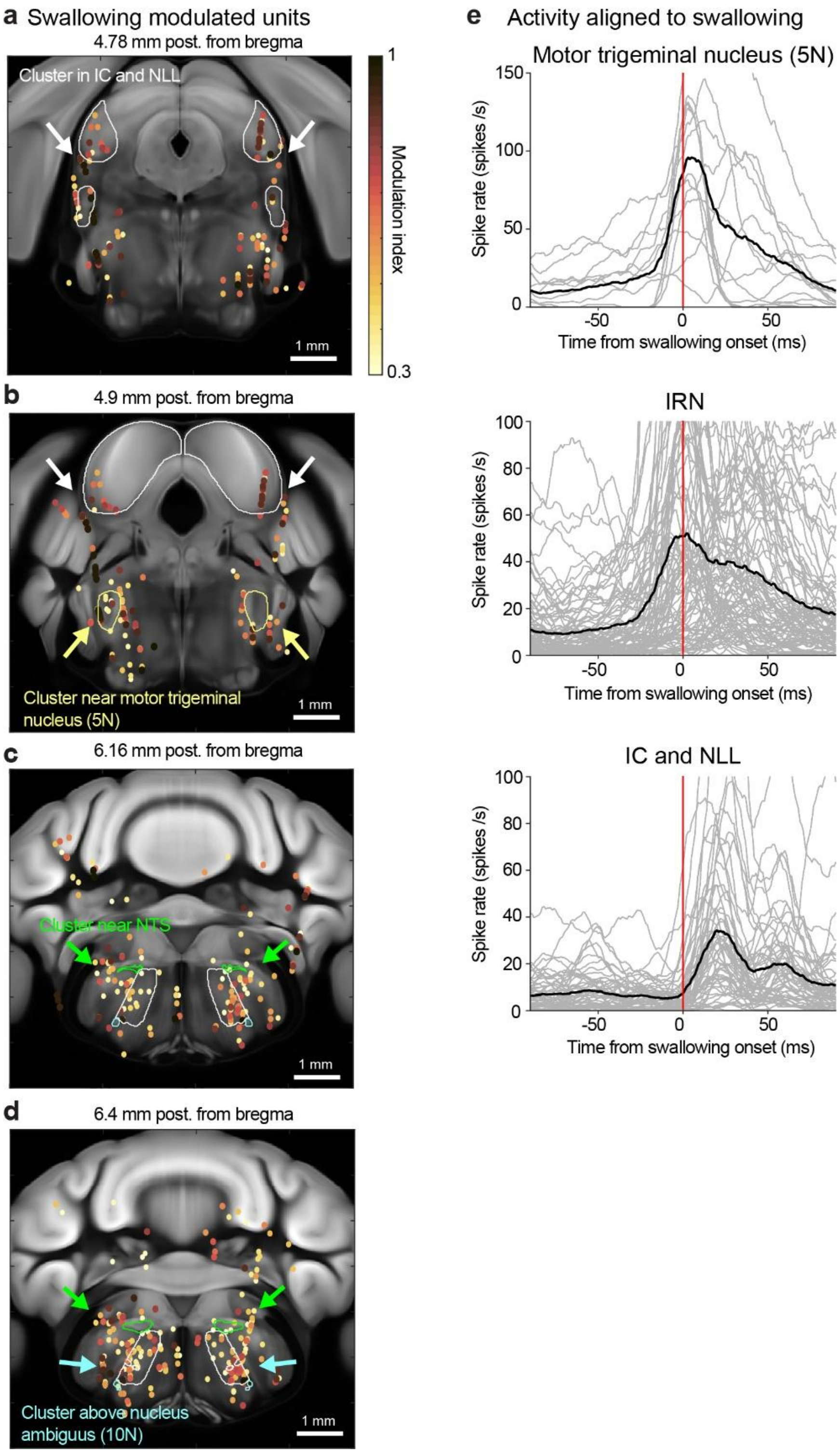


**Extended Data Figure 6. Distribution of the swallowing activity map.**

- **a.** Spatial map of neurons with significant spike rate modulation during swallowing. A cluster of swallowing neurons was observed near the inferior colliculus (IC, white outline) and nucleus of the lateral lemniscus (NLL, white outline).
- **b.** A cluster of swallowing neurons was observed near the motor trigeminal nucleus (5N, yellow outline).
- **c.** A cluster of swallowing neurons was observed near the nucleus of the solitary tract (NTS, green outline) and dorsal IRN (white outline).

**d.** A cluster of swallowing neurons was observed above the nucleus ambiguus (10N, cyan outline) and ventral IRN (white outline).

**e.** Spike rates of swallowing neurons aligned to swallowing onset. Activities in 5N and IRN lead swallowing while activities in the IC and NLL follow swallowing. Gray lines, individual neurons. Black line, mean.

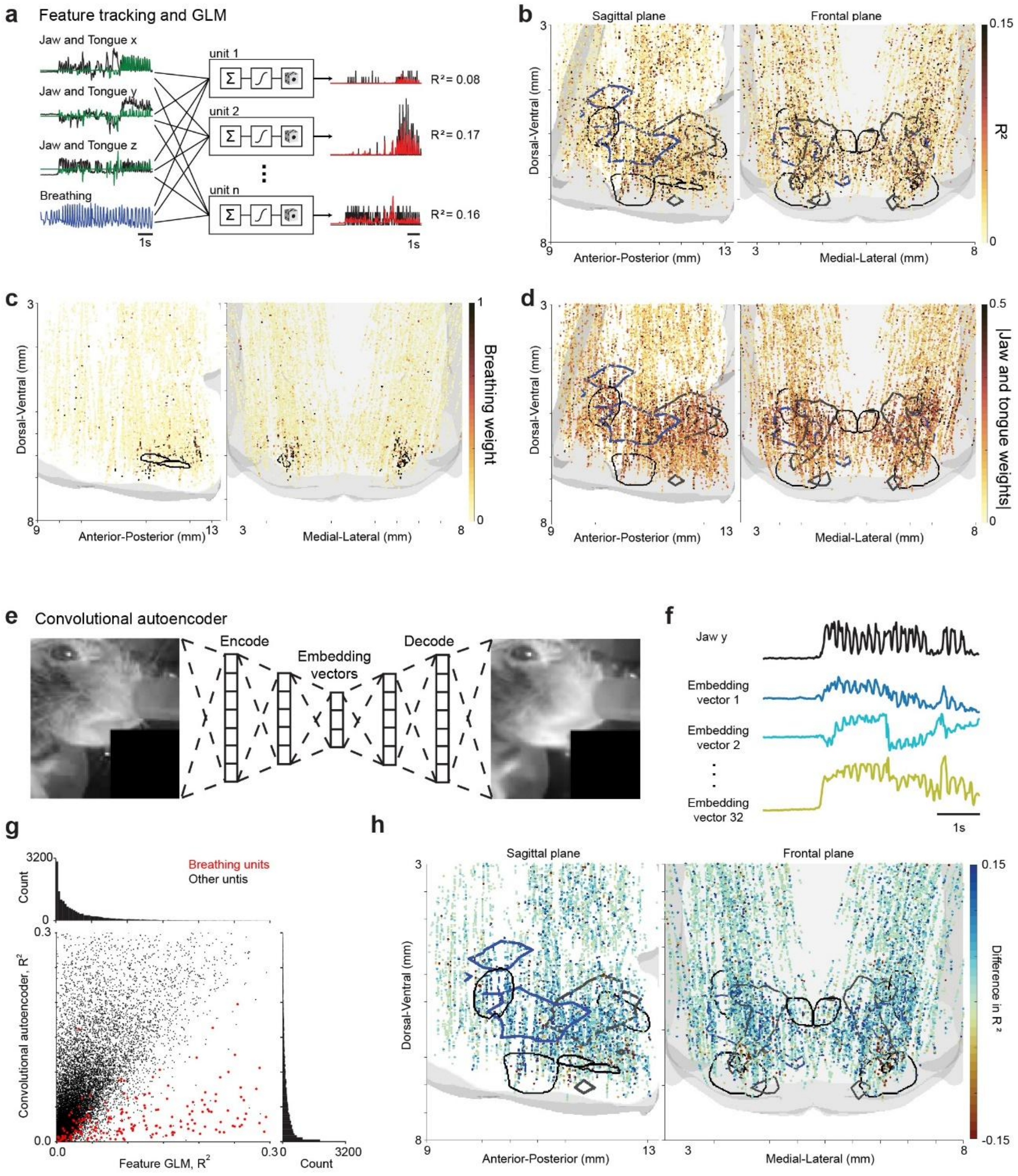


**Extended Data Figure 7. Movement encodings of brainstem units.**

- **a.** Generalized Linear Model (GLM) that uses tracked features of licking and breathing to predict the spike rate of units. The tracked features are the jaw (black) and tongue (green) positions in three dimensions and the airflow measurement (blue). The GLM has a logistic link function followed by Poisson spike statistics. The spike rate of units is shown in black. The predicted spike rate from the GLM is shown in red.
- **b.** Variance account for ($r^2$) of GLM fits for all units. The contours denote the boundaries of motor and premotor nuclei for licking and breathing.
- **c.** Weights of the breathing regressor. The contours denote the boundaries for nucleus ambiguus, 10N, which is above the preBötzinger complex and below the IRN.
- **d.** Mean weights of the jaw and tongue regressors. The contours denote the boundaries of motor nuclei and premotor regions for tongue and jaw muscles.
- **e.** Convolutional Autoencoder (CAE) to capture movements in the video beyond the tracked features. Each frame of the video is downsampled to 120 pixels X 112 pixels as the input layer. The CAE consists of 2 blocks of 3 convolution layers. The embedding is a 32-dimensional vector for each frame.
- **f.** Example embeddings compared to the jaw tracking in black.
- **g.** Scatter histogram of the variance accounted for ($r^2$) of GLM fits, the CAE embeddings versus the tracked features. Each dot represents a unit. Red, breathing-related units.
- **h.** The difference between the $r^2$ of GLM fits, the CAE embedding versus the tracked features. The difference in $r^2$ is plotted against the boundaries of motor and premotor nuclei for tongue and jaw. The $r^2$ of the CAE embedding is higher in most regions of the brainstem but lower in the breathing oscillator.

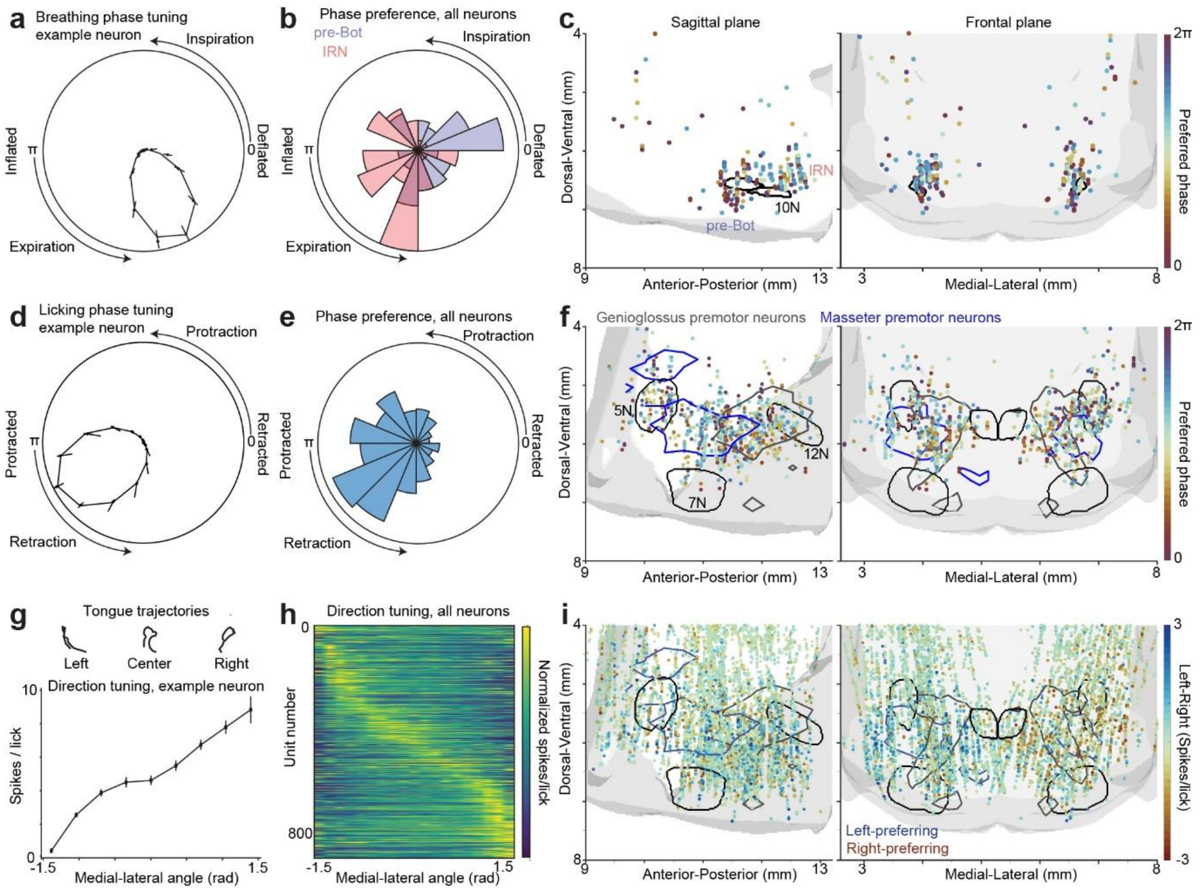


**Extended Data Figure 8. Neural coding of breathing phase, licking phase, and licking direction.**

- **a.** Example breathing-related unit. Error bars represent 95% confidence interval from 100 permutations of the spike distribution.
- **b.** Population histogram of the preferred breathing phase for all the breathing tuned units (Modulation Index > 0.9 and P < 0.001 from the Wilcoxon rank sum test against resampled distribution).
- **c.** Map of preferred breathing phase for all the breathing tuned units plotted against the nucleus ambiguus, 10N, which is above the preBötzinger complex and below the IRN.
- **d.** Example licking-related unit. Error bars represent 95% confidence interval from 100 permutations of the spike distribution.
- **e.** Population histogram of preferred licking phase for all the licking tuned units (Modulation Index > 0.9 and P < 0.001 from the Wilcoxon rank sum test against resampled distribution).
- **f.** Map of preferred licking phase for all the licking tuned units plotted against the motor nuclei and premotor regions for tongue and jaw.
- **g.** Top, tongue tracking from three example licks. The angle of each lick is calculated as the angle between the tip of the tongue at the fully protracted state and the midline. Bottom, example lick direction tuning curve for a licking-related unit. The spikes between -50ms to 100ms relative to the onset of each lick are binned according to the angle of each lick. Error bars represent s.e.m.
- **h.** Lick direction tuning curves for all licking-related units ranked by the cross-validated preferred direction. Each row represents the normalized tuning curve of a unit.
- **i.** Activity map of the spike rate of left licks subtracting the spike rate of right licks for all units. Activity in each hemisphere is selective for ipsilateral lick direction.

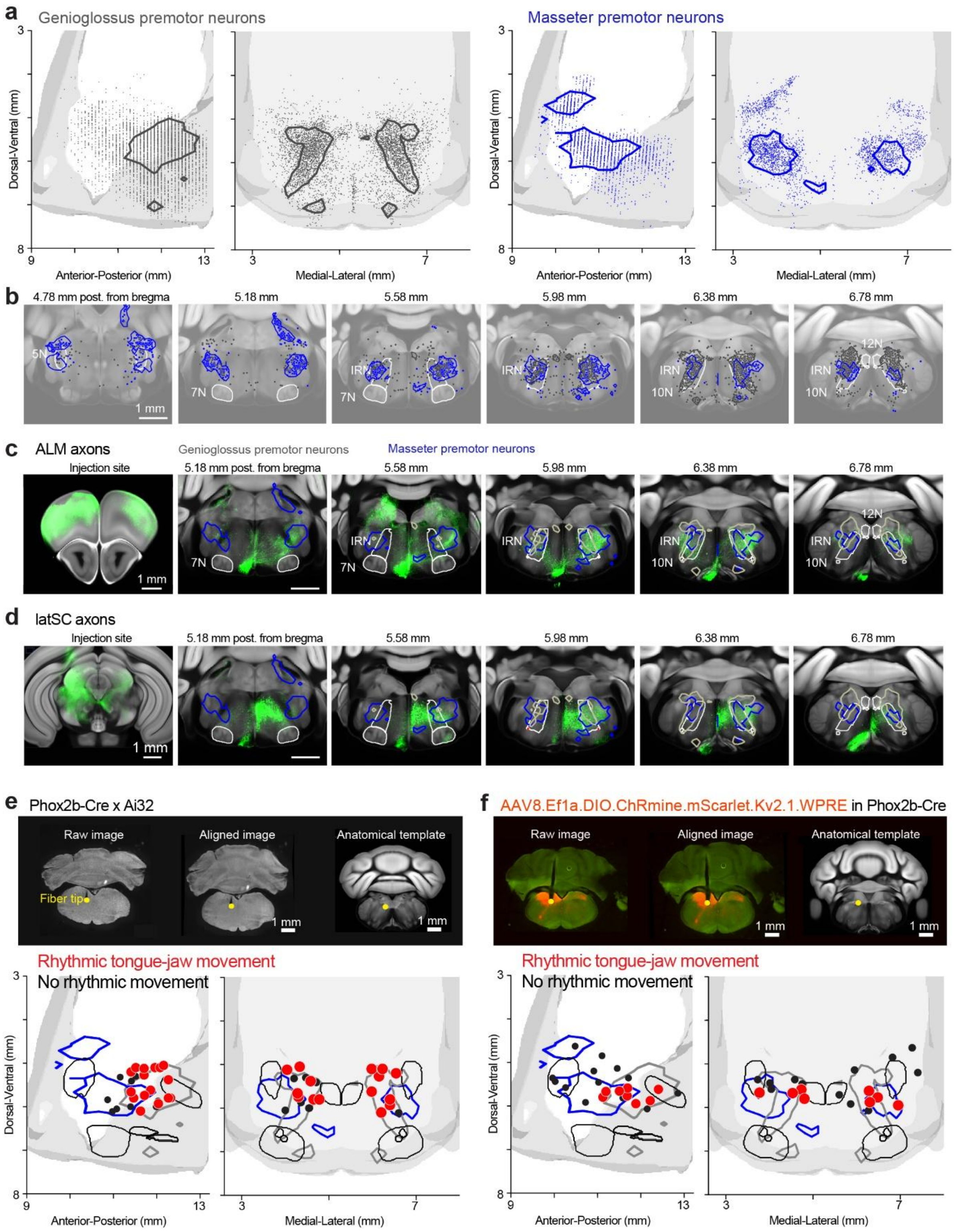


**Extended Data Figure 9. Analysis of anatomy and photostimulation sites to map brainstem premotor network for licking.**

- **a.** Premotor neurons of the genioglossus (gray dots) and masseter muscles (blue dots). The gray boundary denotes a density of 0.0063 neurons/mm$^3$. The blue boundary denotes a density of 0.0019 neurons/mm$^3$.
- **b.** Coronal sections showing the premotor neurons of the genioglossus and masseter muscles.

**c.** Fluorescence from ALM axon projections in green overlaid on the contours of the premotor neurons as well as the brainstem nuclei involved in orofacial movements, 7N: Facial Nucleus. 10N: Nucleus Ambiguus. 12N: Hypoglossal Nucleus. IRN: Intermediate Reticular Nucleus.

**d.** Fluorescence from latSC axon projections. Same as **c**.

**e.** *Top*, example photostimulation sites in Phox2b-Cre x Ai32 mice showing the optical fiber tip location and alignment to CCF. *Bottom*, spatial map of photostimulation sites in Phox2b-Cre x Ai32 mice. Red dots, photostimulation sites where rhythmic licking are elicited. Black dots, photostimulation sites where rhythmic licking are not elicited. N=25 photostimulation sites, 16 mice.

**f.** *Top*, example photostimulation sites in Phox2b-Cre mice showing ChRmine-mScarlet expression and alignment to CCF. *Bottom*, spatial map of photostimulation sites in Phox2b-Cre mice. Same as **d**. N=21 photostimulation sites, 11 mice.

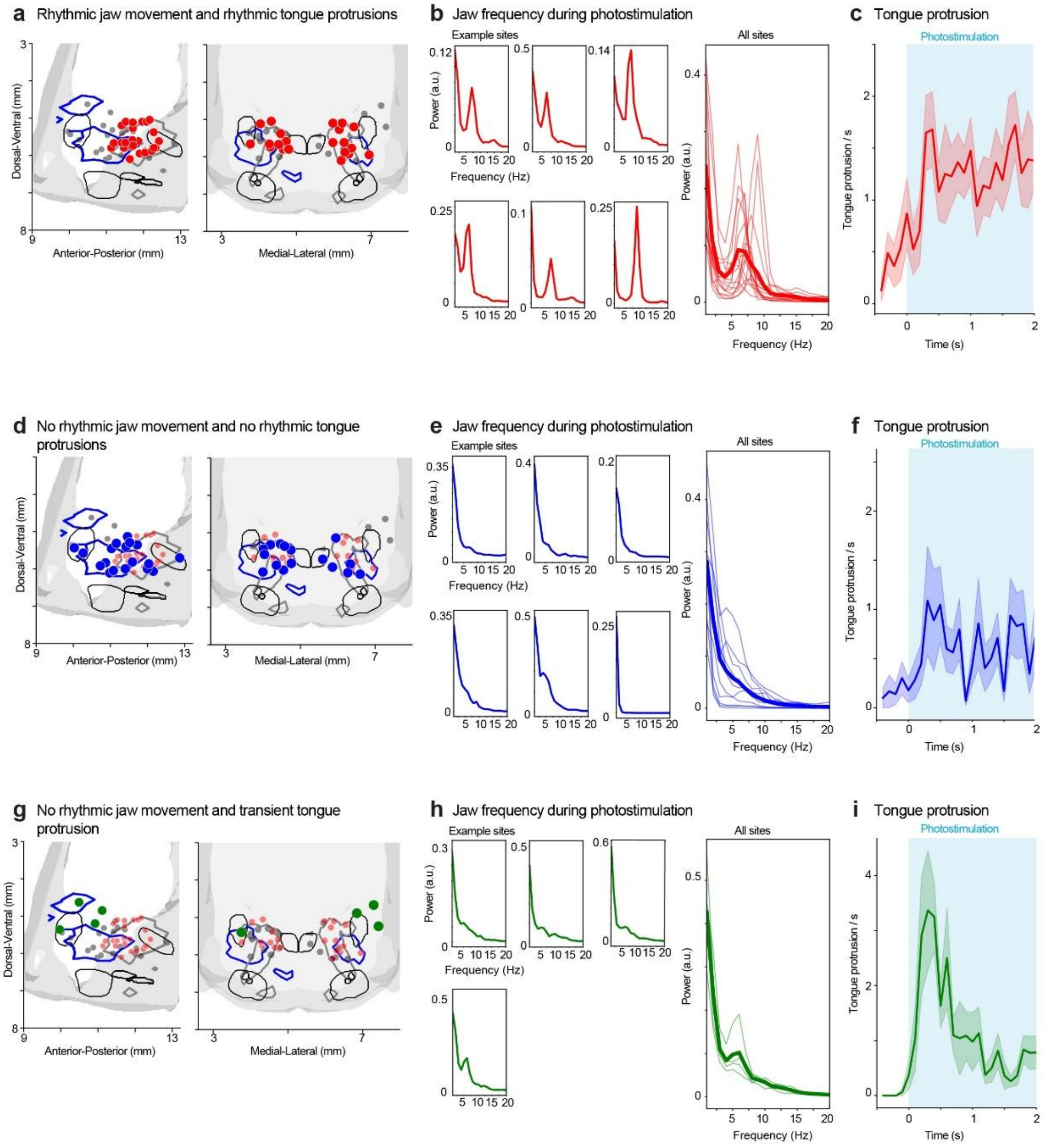


**Extended Data Figure 10. Analysis of tongue and jaw movements evoked by brainstem photostimulation.**

- **a.** Spatial map of photostimulation sites where rhythmic jaw movement and rhythmic tongue protrusions are evoked by constant photostimulation.
- **b.** The rhythmicity of jaw movement is quantified using Fourier transform. *Left,* jaw movement frequency of individual photostimlation sites. Right, all photostimulation sites, N=18. Thin lines, individual sites; thick lines, average. A distinctive peak is observed at 7 Hz, which is the natural frequency for licking (Fig 1d).
- **c.** Tongue protrusion evoked by photostimulation. Mean ± s.e.m. across photostimulation sites. Rhythmic tongue protrusions are evoked during the entire photostimulation epoch.
- **d.** Spatial map of photostimulation sites where no rhythmic jaw movement are evoked by photostimulation.
- **e.** Same as **b**, but for photostimulation sites in **d**. N=11.
- **f.** Same as **c**, but for photostimulation sites in **d**.
- **g.** Spatial map of photostimulation sites where no rhythmic jaw movement are evoked but photostimulation evoked a transient tongue protrusion at the beginning of photostimulation. The tongue protrusion stopped after 1 or 2 licks.
- **h.** Same as **b**, but for photostimulation sites in **g**. N=4.
- **i.** Same as **c**, but for photostimulation sites in **g**.

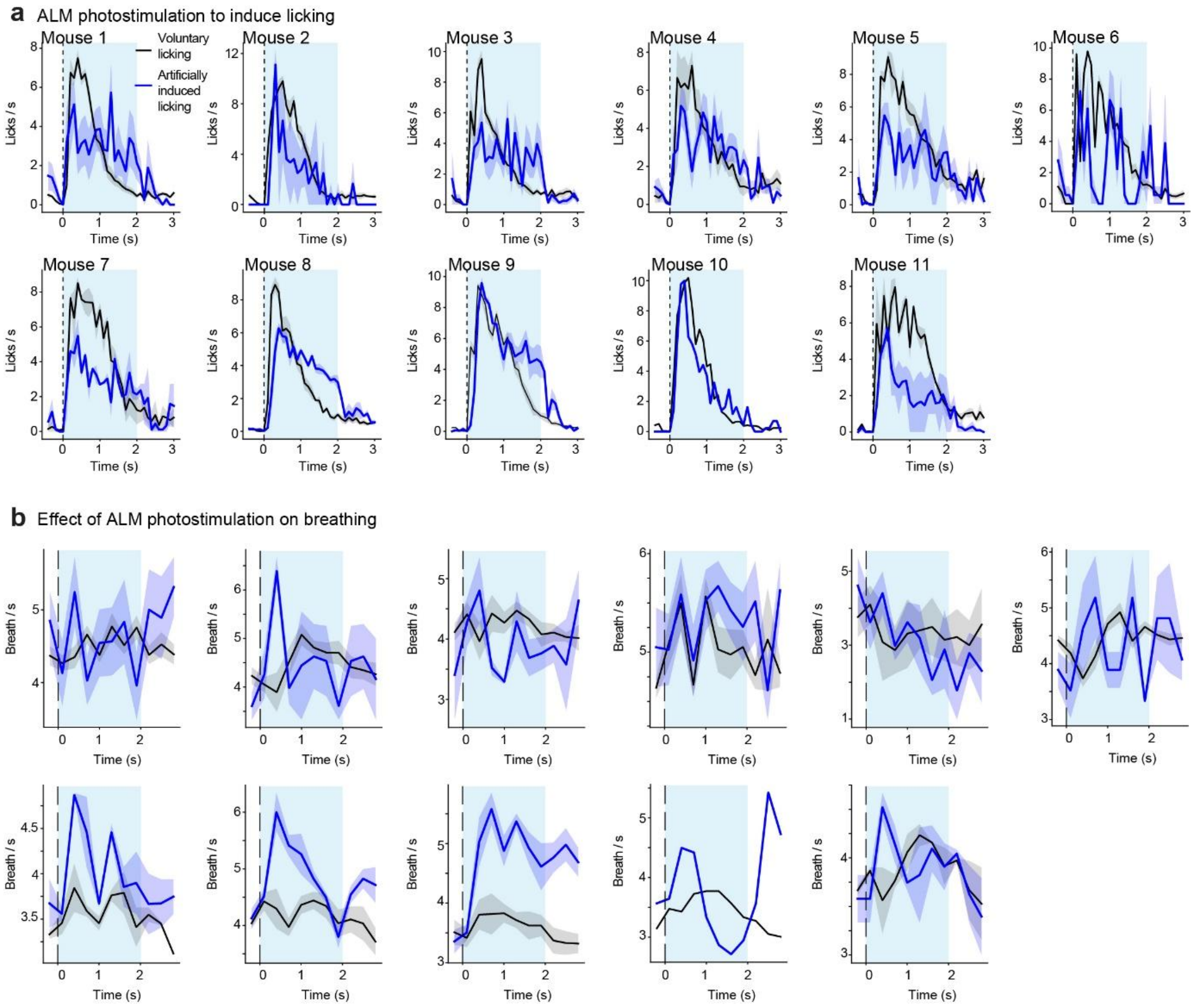


**Extended Data Figure 11. Analysis of licking and breathing during ALM photostimulation.**

- **a.** Lick rate for individual mice in the cued licking task. Licking (black) is aligned to the Go cue (dashed line). Artificially induced licking (blue) is aligned to the photostimulation (cyan). Mean ± s.e.m. across session. For mouse 10, only a single session was tested so only the mean is shown.
- **b.** Breathing rate for individual mice in the cued licking task. Same as **a**.

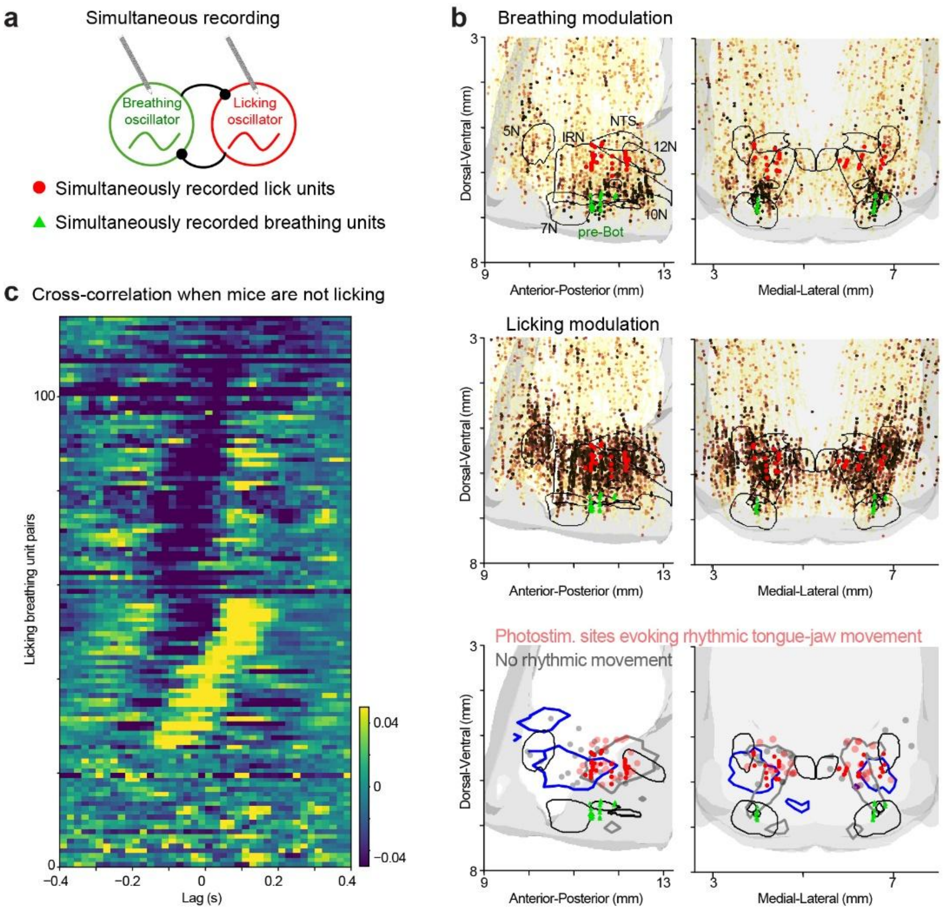


**Extended Data Figure 12. Simultaneous recording of breathing and licking oscillators.**

- **a.** Schematic. Only simultaneously recorded breathing and licking units near the preBötzinger complex and posterior-dorsal IRN/PARN (putative licking oscillator) are selected for analysis.
- **b.** Spatial maps of the simultaneously recorded breathing units (red circles) and licking units (green triangles). *Top,* units overlaid on the breathing activity map, data from Fig 2f. *Middle,* units overlaid on the licking activity map, data from Fig 2g. *Bottom,* units overlaid on the photostimulation map where constant stimulation can drive sustained rhythmic licking (faint red dots), data from Fig 3h.
- **c.** Cross-correlogram of the unit pairs during the baseline period of the cued licking task when mice were not licking. Same as Fig 4g.
